# Towards community-driven visual proteomics with large-scale cryo-electron tomography of *Chlamydomonas reinhardtii*

**DOI:** 10.1101/2024.12.28.630444

**Authors:** Ron Kelley, Sagar Khavnekar, Ricardo D. Righetto, Jessica Heebner, Martin Obr, Xianjun Zhang, Saikat Chakraborty, Grigory Tagiltsev, Alicia K. Michael, Sofie van Dorst, Florent Waltz, Caitlyn L. McCafferty, Lorenz Lamm, Simon Zufferey, Philippe Van der Stappen, Hugo van den Hoek, Wojciech Wietrzynski, Pavol Harar, William Wan, John A.G. Briggs, Jürgen M. Plitzko, Benjamin D. Engel, Abhay Kotecha

## Abstract

*In situ* cryo-electron tomography (cryo-ET) has emerged as the method of choice to investigate structures of biomolecules in their native context. However, challenges remain in the efficient production of large-scale cryo-ET datasets, as well as the community sharing of this information-rich data. Here, we applied a cryogenic plasma-based focused ion beam (cryo-PFIB) instrument for high-throughput milling of the green alga *Chlamydomonas reinhardtii*, a useful model organism for *in situ* visualization of numerous fundamental cellular processes. Combining cryo-PFIB sample preparation with recent advances in cryo-ET data acquisition and processing, we generated a dataset of 1829 reconstructed and annotated tomograms, which we provide as a community resource to drive method development and inspire biological discovery. To assay the quality of this dataset, we performed subtomogram averaging (STA) of both soluble and membrane-bound complexes ranging in size from >3 MDa to ∼200 kDa, including 80S ribosomes, Rubisco, nucleosomes, microtubules, clathrin, photosystem II, and mitochondrial ATP synthase. The majority of these density maps reached sub-nanometer resolution, demonstrating the potential of this *C. reinhardtii* dataset, as well as the promise of modern cryo-ET workflows and open data sharing towards visual proteomics.

## Introduction

The concept of “visual proteomics” was first proposed nearly two decades ago, before instrumentation was developed to enable it^1^. At its heart is the idea of using cryogenic electron tomography (cryo-ET) to produce comprehensive structural inventories of macromolecular complexes inside native cells (*in situ*). The term visual proteomics has since taken on broader interpretations, reflecting the growing potential of integrating cryo-ET with diverse methodologies such as mass spectrometry and AI-based structure prediction^2^. Such efforts are limited by the availability and quality of cryo-ET data, but thanks to advances in instrumentation and computation, cryo-ET is now entering a renaissance where the dream of visual proteomics is beginning to become tangible^3,4^. However, this ambitious goal requires continued developments in 1) data collection to extensively sample cellular environments with high resolution, 2) data analysis to identify and structurally characterize diverse species of macromolecules within these environments, and 3) data sharing in community repositories that enable biological exploration, method development, and training of neural networks.

The quality and throughput of cryo-ET data acquisition has seen substantial improvements in recent years. A crucial sample preparation step for cryo-ET is the thinning of vitreous biological material, most commonly performed with a focus ion beam (FIB) to produce cellular “lamellae” with a thickness of ∼100-300 nm. Advances in cryo-FIB milling of plunge-frozen and high-pressure frozen biological material have enabled exploration of diverse cell types^5–12^ and even multicellular tissue^13–15^. Throughput has increased by an order of magnitude as a result of automated FIB milling procedures^16–19^ and multishot cryo-ET, which allows multiple tomograms to be acquired in parallel^20–24^. Despite these gains, the current generation of gallium-based FIB instruments are still limited by manual loading of only two grids at a time, slow milling speeds, and ice contamination on the milled lamella surfaces. A new generation of plasma-based FIB (PFIB) instruments promises to improve the speed and quality of lamella production^25^. Unlike gallium beams that tend to diverge at high beam currents, plasma sources produce a more tightly focused probe at high beam currents^26^, thereby facilitating efficient milling of samples ranging from small cells to large volumes of multicellular tissue. In addition, some modern cryo-PFIB systems are equipped with robotic sample handling and automated liquid nitrogen filling that further increase throughput and enable automated milling sessions over long runtimes^25^.

As cryo-ET data output increases, visual proteomics will rely on computational developments aimed at extracting the wealth of information in cellular tomograms. Advances in template matching^27–31^ and subtomogram averaging (STA)^29,32,33^ have recently enabled the determination of sub-nanometer-resolution structures inside cells for complexes that are relatively large and abundant^34–37^. So far, the most popular targets for *in situ* structural analysis have been ribosomes, which have reached near-atomic resolution (3-4 Å) and enabled classification of numerous native conformational states^38–43^. Alongside these gains in native structure determination, there has been rapid development in automated tomogram segmentation and particle detection within cellular volumes, enabled by increasingly powerful neural networks^44–48^.

However, these AI-based annotation approaches and *in situ* STA both require large and diverse cryo-ET datasets for robust training of generalized neural networks and identifying enough examples of less abundant macromolecular species to allow high-resolution averaging. Public data repositories, such as the Electron Microscopy Public Image Archive (EMPIAR)^49^ and the Chan Zuckerberg Imaging Institute (CZII) CryoET Data Portal^50^, aim to aggregate cryo-ET datasets from the global research community to enable method development and biological discovery. Yet to date, very few large-scale datasets of cellular cryo-ET are available.

Here, we provide a new resource to enable community-driven efforts towards visual proteomics. Using an efficient workflow that combines cryo-PFIB milling with high-resolution cryo-ET, we produced an open community dataset of 1829 curated tomograms (and accompanying raw data) covering the breadth of organelles found within *Chlamydomonas reinhardtii* cells. These unicellular green algae are ideal models for cryo-ET, thanks to their small size (compatible with plunge-freezing vitrification and high-throughput FIB milling), good cryo-ET contrast, and reproducible cellular architecture, with a cornucopia of eukaryotic structural cell biology to explore^51^. Furthermore, *C. reinhardtii* benefits from a wealth of molecular tools and mutant libraries^52,53^, as well as an active research community using this model organism to study diverse questions including cilia biology and photosynthesis^54–56^. To test the feasibility of structure determination from our dataset, we performed STA of a variety of native macromolecules ranging in size from ∼200 kD to >3 MDa, including both soluble and membrane-integral complexes. It is our hope that this publicly available *C. reinhardtii* dataset will empower new discoveries and method development, while providing an example for open sharing of future large-scale datasets generated by the rapidly growing cryo-ET community.

## Results

### Cryo-PFIB milling to enable large-scale cryo-ET

We leveraged the automated features of the recently developed Arctis cryo-PFIB to increase the throughput of lamella production. Each session, we used the autoloader and multi-specimen cassette to screen 8-10 grids with vitrified *C. reinhardtii* cells before beginning milling. The suitability of each grid was assessed with scanning electron microscopy (SEM) atlas overviews (Figs. 1A, S1A). Grids with sufficient coverage of isolated single cells or clusters of up to three cells, close to the center of the grid square and with minimal vitrified buffer surrounding the cells, were chosen for further processing. Grids that had larger clumps of cells or a thicker ice layer were discarded, as such material is unlikely to be completely vitrified.

**Figure 1.**
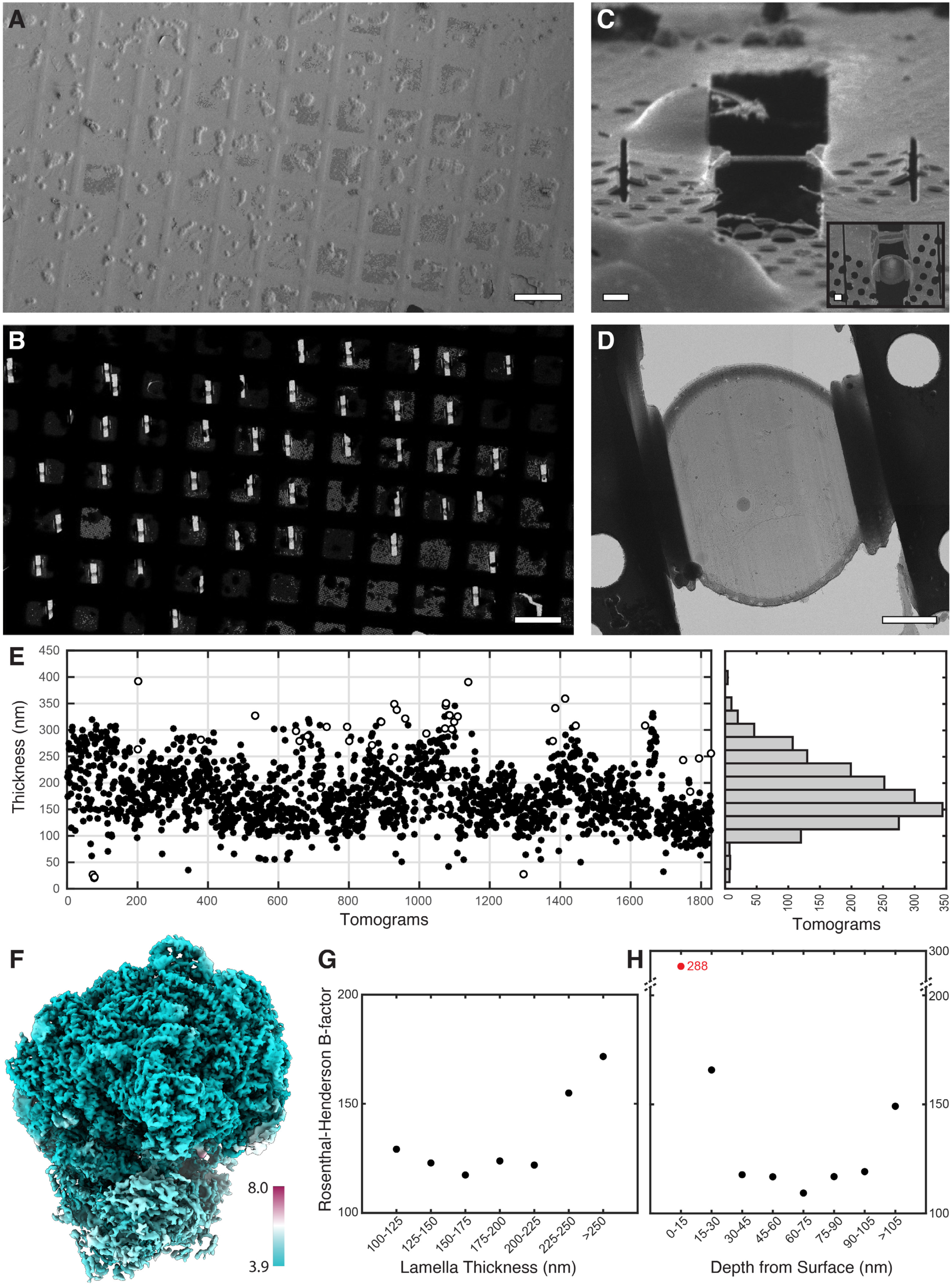
High throughput cryo-ET workflow applied to *C. reinhardtii*. **A)** SEM image of an EM grid covered in frozen *C. reinhardtii* cells. **B)** Low magnification TEM image of the grid with lamellae that were prepared by cryo-PFIB milling. **C)** Ion beam image of a single lamella as seen from the milling angle. Inset: SEM top view of the lamella. **D)** TEM overview search map of a single lamella. **E)** Plot of tomogram thickness for the full dataset (EMPIAR-11830), summed into the histogram on the right. Tomogram thickness determined by two methods (black circles: TOMOMAN functionality that uses IMOD’s findsection program, white circles: Slabify; see methods). **F)** Local-resolution display of the 80S ribosome subtomogram average, with the large subunit at Nyquist resolution (3.92 Å). **G)** Dependence of 80S ribosome B-factor on lamella thickness. **H)** Dependence of 80S ribosome B-factor on the distance of particles from the lamella surface. Note that the 0-15 nm bin (red point) contains partial ribosomes that were cut by FIB milling, thus resulting in a much higher B-factor. The 15-30 nm bin contains complete ribosomes but a higher B-factor than deeper bins spanning 30-105 nm, likely due to beam damage near the lamella surface. Scale bars: 100 µm in A and B; 1µm in C, C inset, and D.

Our PFIB system was equipped with xenon, argon, and oxygen ion sources, each offering distinct properties. Characterization of milling with different ion species has shown some differences^57^, including higher sputter rates when milling with xenon. While cryo-FIB/SEM volume imaging and lamella preparation have traditionally utilized either gallium or argon ion beams^16,25,58–60^, a recent study demonstrated the successful use of xenon for milling high-pressure frozen biological samples^61^. In this work, the majority of lamellae were milled with xenon, but we also milled a smaller set of lamellae using argon to compare the milling throughput. The average time to prepare a lamella from *C. reinhardtii* cells with xenon ions was ∼23 min, including 4 min site preparation time and 19 min milling time (Table 1). Lamellae prepared with argon ions averaged ∼26 min per site, with 4 min site preparation time and 22 min milling time (Table 2). Overall, we found that xenon was marginally faster for milling lamellae from plunge-frozen *C. reinhardtii* cells. Depending on the number of available cells, between 10 and 40 lamellae were prepared with an automated milling routine on each grid (Figure 1B-D) using AutoTEM^19^ or WebUI^62^ software. The target width of each lamella was 8-10 μm, and the target thickness was 150-200 nm.

**Table 1.** Milling parameters for xenon ions on *C. reinhardtii* cell (8 µm wide lamellae)

| **Milling steps** | **Milling current (pA)** | **~ Milling time** |
| --- | --- | --- |
| Stress relief cut | 100 | 1.5 min |
| Rough milling | 300 | 5 min x2 |
| Medium milling | 100 | 1.5 min x2 |
| Fine milling | 100 | 55s x2 |
| Polishing | 30 | 1 min 15 s x2 |

**Table 2.** Milling parameters for argon ions on *C. reinhardtii* cell (8 µm wide lamellae)

| **Milling steps** | **Milling current (pA)** | **~ Milling time** |
| --- | --- | --- |
| Stress relief cut | 200 | 1 min |
| Rough milling | 740 | 5 min x2 |
| Medium milling | 200 | 1.9 min x2 |
| Fine milling | 200 | 1.2 min x2 |
| Polishing | 20 | 2.2 min x2 |

Grids with lamellae were transferred to the Krios transmission electron microscope (TEM) directly from the Arctis cryo-PFIB using the same cassette and capsule setup, eliminating manual handling steps and, thereby, reducing the risks of ice contamination and lamella breakage during the transfer process. This direct transfer prevents grid rotation between the two instruments, ensuring that lamellae are in the correct orientation for tilt-series collection, with the FIB milling direction roughly perpendicular to the TEM tilt axis (Fig. S1B-C). Across 42 overnight imaging sessions, we acquired a dataset of 2991 tilt-series from 55 different grids containing over 300 lamellae. As a tradeoff between cellular coverage and resolution, we selected a pixel size of 1.96 Å, setting the Nyquist limit at 3.92 Å. All tilt-series and their individual tilt images were manually curated and then processed through an automated alignment and reconstruction pipeline. Tilt-series that suffered from non-vitreous ice and alignment issues were discarded. In total, 1829 reconstructed tomograms were deemed to be of good quality for further analysis. The high discard rate was primarily the result of inconsistent sample quality, with some grids containing mostly non-vitreous cells. We measured the thicknesses of all the retained tomograms, which varied from ∼100 to ∼300 nm (excluding ∼10% outliers), with a median thickness of 157 nm (Fig. 1E). To aid visual inspection, segmentation, and particle picking, all tomograms were denoised using cryo-CARE^63^.

### Analysis of radiation damage from PFIB using 80S ribosomes

Recently, sub-5 Å STA maps of 80S ribosomes were obtained from cryo-ET data acquired on cells milled using cryo-PFIB milling^25,64^. Several functional states of the 80S ribosome complex were resolved from these datasets using classical^29^ as well as machine learning-based^64^ approaches. Here, we used *C. reinhardtii* 80S ribosome particles to assess the dependence of attainable resolution on lamella thickness and PFIB-induced beam damage to the lamella surface. We first used a small-subset of tomograms collected from a single lamella. With ∼6,000 subvolumes extracted from 16 reconstructed tomograms, we obtained a ribosome structure with a global resolution of 5.5 Å (Fig. S2B). We then expanded our dataset to ∼140K subvolumes from 600 tomograms and obtained a 4.0 Å STA map, with resolution of the ribosomal large subunit extending to the Nyquist limit (3.9 Å) (Figs. 1F, S2D). In addition, the large size of the dataset allowed us to use subvolume classification (Fig. S2E) to distinguish free cytosolic ribosomes (in complex with ES27 and Arx1) from ribosomes bound to the ER membrane (in complex with sec61-TRAP) (Fig. S2F)^65^, as well as different functional states with resolved occupancies for tRNA and elongation factors (Fig. S2G).

Recent studies have demonstrated that gallium and argon ion beams damage the lamella surface during cryo-FIB milling^25,66,67^. We analyzed Rosenthal-Henderson B-factors for 80S ribosome averages generated from lamellae with varying thicknesses and observed that thicker lamellae (>225 nm) had a reduced signal-to-noise ratio (SNR) (Figs. 1G, S3). Although this suggests that it is beneficial to mill thinner lamellae for high-resolution imaging, further B-factor analysis for ribosomes at varying depth within lamellae indicated a damage layer extending up to 30 nm from the lamella surface (Figs. 1H, S4), consistent with previous reports^25,67^. It’s important to note that this analysis has inherent limitations due to the size of 80S ribosomes, which are ∼30 nm in diameter. Therefore, the high B-factor value for particles 0-15 nm from the surface (B-f = 288) is due to incomplete ribosomes that were cut by FIB milling. However, the 15-30 nm bin has a higher B-factor (B-f = 166) than the 30-45 nm bin (B-f = 118), indicating that indeed particles 15-30 nm from the lamella surface incur some damage. The B-factors remain relatively constant between 30-105 nm, but increase again in the 105+ nm bin likely due to reduced SNR in thicker lamellae. From this analysis, we conclude that *C. reinhardtii* lamellae of <225 nm thickness are best for high-resolution STA, using particles positioned approximately 30-100 nm from the lamella surface.

### Towards visual proteomics in Chlamydomonas

*C. reinhardtii* has characteristic cellular architecture, which has enabled *in situ* cryo-ET studies of the nucleus (including nucleolus and nuclear pores)^68–70^, endoplasmic reticulum^65,71^, Golgi and secretory system^72–74^, centrioles and cilia^75–79^, actin^80^, mitochondria^81^, and chloroplasts (including pyrenoid and thylakoids)^82–85^. To gain an overview of this cellular architecture, we first used a Hydra cryo-PFIB to perform slice-and-view volume EM imaging of an entire vitreous *C. reinhardtii* cell (Fig. 2A, S5). This method, traditionally established for resin-embedded biological samples^86,87^ and recently demonstrated for cryo applications^58,60^, works by sequentially removing material with the ion beam and imaging the newly created cross-section surfaces with the scanning electron beam to obtain a 3D volume. The resulting overview of a full *C. reinhardtii* cell helped provide orientation when examining low-magnification search images of lamellae from other cells in the TEM (Fig. 2B), guiding the selection of positions for tilt-series acquisition. The 1829 tomograms in the curated dataset contain the following organelles (sometimes with multiple organelles in one tomogram): 387 tomograms of the nucleus (Fig. 2C), 246 of the Golgi apparatus (Fig. 2D), 16 of centrioles (Fig. 2E), 581 of mitochondria (Fig. 2F), and 823 of the chloroplast (Fig. 2G), including 84 of the pyrenoid compartment (Fig. 2H). In addition, microtubules, actin, endoplasmic reticulum (ER), and a variety of acidocalcisomes and other vacuoles are also present in the dataset. We trained several 2D, 2.5D, and 3D U-nets on a small subset of the data to enable semi-automated semantic segmentation of this cellular ultrastructure^45,88,89^ (Figs. 2C-H, S6).

**Figure 2.**
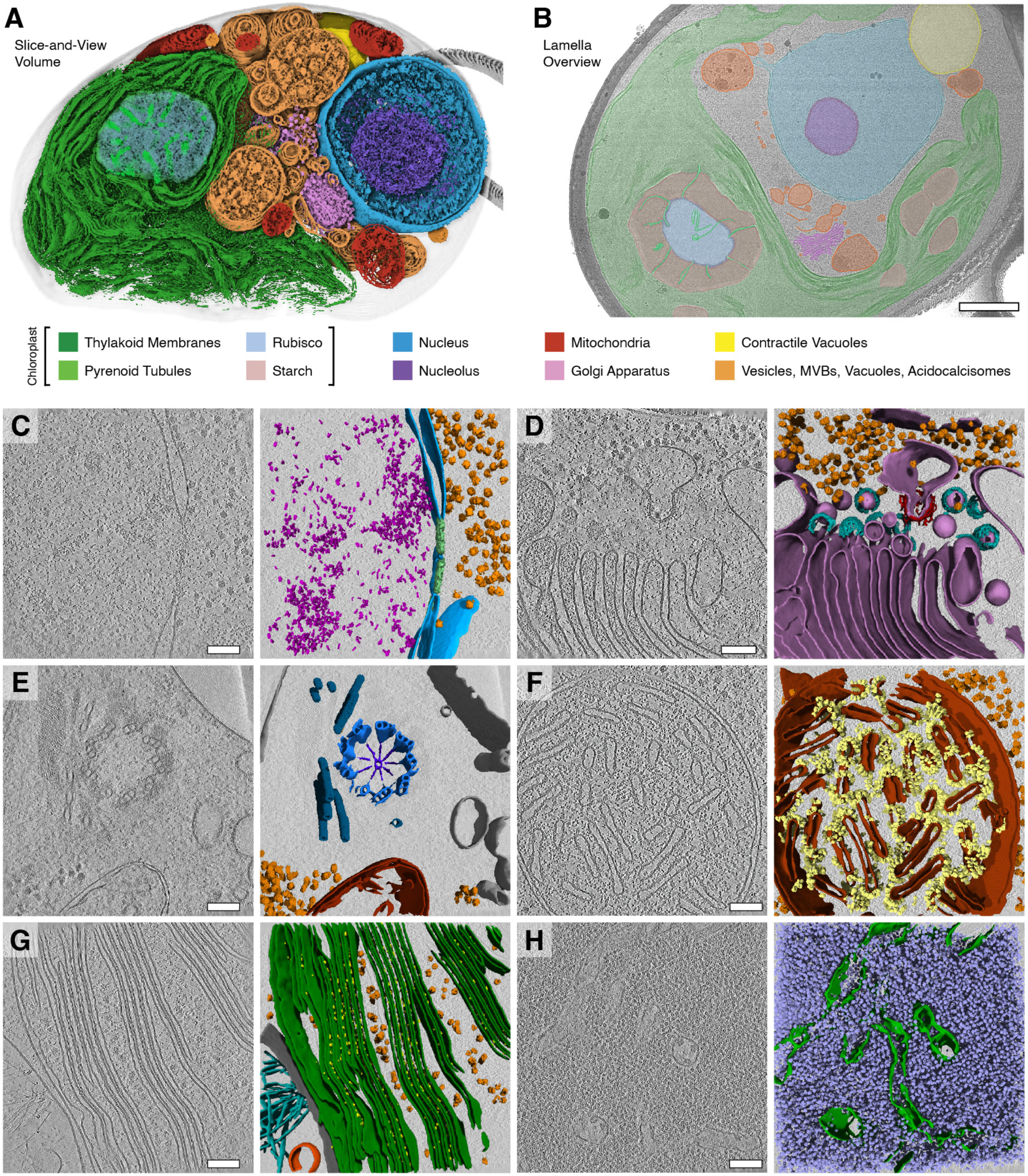
Molecular architecture of *C. reinhardtii* cellular compartments found in the large-scale cryo-ET dataset. **A)** 3D segmentation of a whole vitreous *Chlamydomonas* cell imaged by cryo-PFIB/SEM slice-and-view (EMPIAR-11275) highlighting overall cellular architecture. 20 nm slice thickness, 335 slices covering about 6.7 μm^3^ of cell volume. **B)** TEM search image of a lamella with major structures identified. Scale bar: 1μm. Color legend applies to panels A and B. **C-H)** Slices through representative tomograms of cellular compartments (left) with corresponding 3D segmentations (right). Scale bars: 100 nm. **C)** Nucleus, with segmented nuclear envelope (blue), nucleosomes (magenta), 80S ribosomes (orange), and a nuclear pore complex (green). **D)** Golgi apparatus, with segmented Golgi and ER membranes (pink), 80S ribosomes (orange), COPII (red) and COPI (cyan). **E)** Basal body, with segmented membranes (grey), mitochondria (red), microtubule triplets (blue), cartwheel (indigo), 80S ribosomes (orange), and rootlet microtubules (navy). **F)** Mitochondria, with segmented mitochondrial membranes (red), ATP synthases (yellow), and 80S ribosomes (orange). **G)** Chloroplast, with segmented thylakoid membranes (green), Photosystem II (yellow), 80S ribosomes (orange), membrane (grey), filaments (cyan), and a vesicle (deep orange). **H)** Pyrenoid, with segmented pyrenoid tubule membranes (green) and Rubisco (lavender blue).

Our deep sampling of *C. reinhardtii* cellular architecture provided views of specific and transient organelle features. For example, we observed a mitochondrion bound to three microtubules via thin filaments (Fig. S6A), as well as a mitochondrion undergoing a putative fission event mediated by contact with the ER (Fig. S6B)^90^. We also acquired tomograms of the ciliary transition zone with assembling intraflagellar transport trains (Fig. S6C)^77,79^, and captured detailed views of the pyrenoid tubules and thylakoid-derived minitubules (Fig. S6D), enigmatic structures that are not fully understood on a molecular level^82,91–93^.

### Subtomogram averaging of diverse molecular complexes

Cytosolic ribosomes have received considerable attention in recent *in situ* cryo-ET studies, as their abundance and large size enables high-resolution structure determination of numerous functional states^38–42^. Indeed, ribosomes were also our first target when benchmarking the quality of our dataset (Figs. 1F-H, S2-S4). However, one of the main aims of producing a large-scale dataset covering a range of *C. reinhardtii* organelles is to enable structural studies of diverse molecular targets within the native cellular environment, taking a step towards visual proteomics. To assess the potential of our dataset, we performed STA of six molecular complexes that varied in size, properties, and cellular location (Fig. 3). To test smaller soluble complexes, we averaged ∼520 kDa Rubisco enzymes in the pyrenoid subcompartment of the chloroplast and ∼200 kDa nucleosomes in the nucleus. To test oligomeric assemblies, we averaged microtubule (MT) filaments and clathrin coats in the cytosol. Finally, to test membrane protein complexes, we averaged ∼600 kDa photosystem II (PSII) complexes in the thylakoid membranes of the chloroplast and ∼1.3 MDa dimers of ATP synthase in the crista membranes of the mitochondria. Five of these averages (all except PSII) reached sub-nanometer resolution, revealing secondary structure features. As detailed in the methods, each of the STA projects was performed by a different user, using a range of computational workflows and software packages for particle picking, alignment, and refinement, thereby demonstrating the utility of the *C. reinhardtii* dataset for a variety of analysis pipelines.

**Figure 3.**
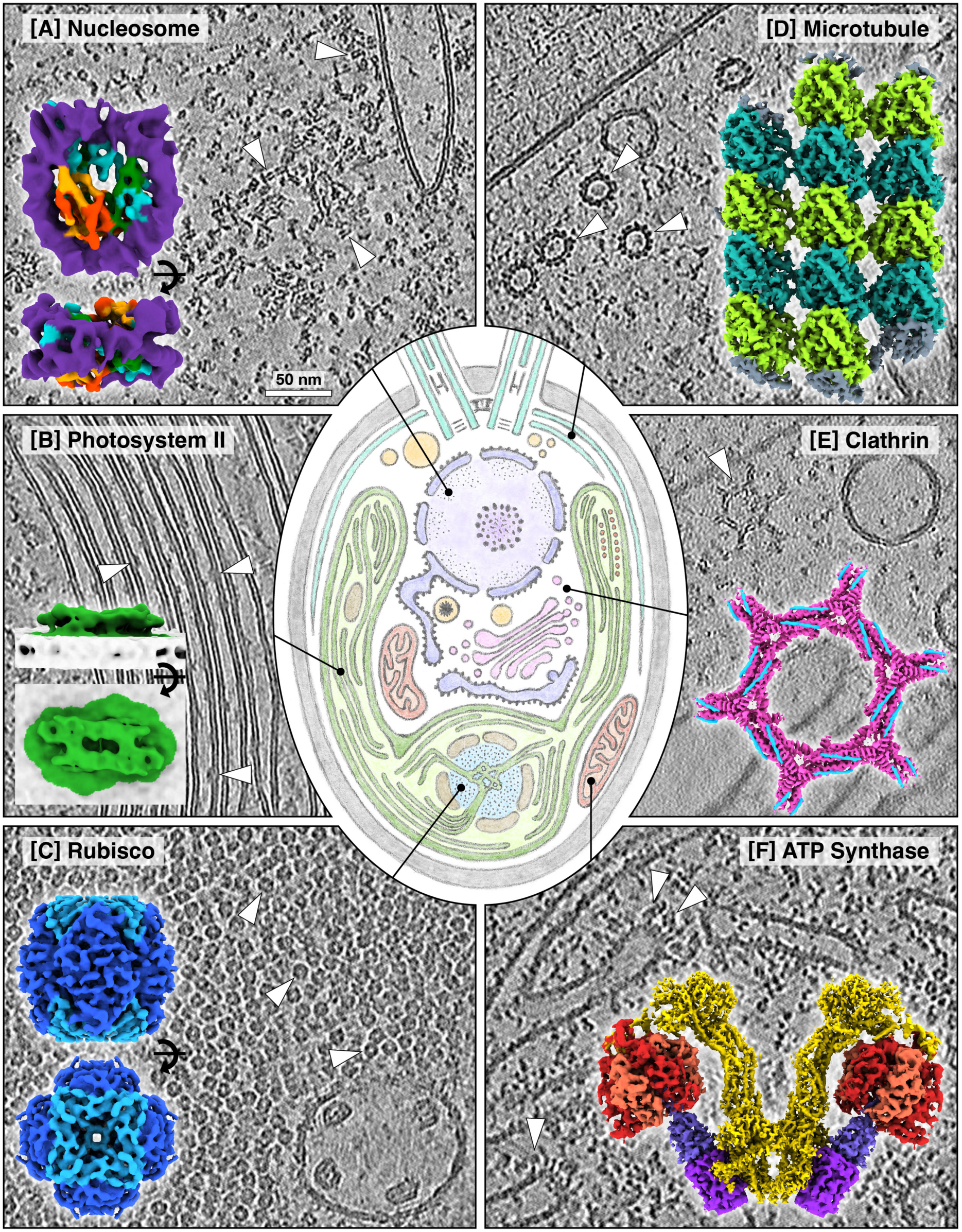
Visualizing molecular machinery inside native *C. reinhardtii* cells. Central illustration: stereotypical cross-section through a Chlamydomonas cell (teal: rootlet microtubules and centrioles; darker purple: nuclear envelope and ER; light purple: nucleoplasm with chromatin at periphery and nucleolus at center; pink: Golgi and associated vesicles; light orange: assorted vacuolar compartments including acidocalcisome with dark granule and contractile vacuoles near centrioles; darker green: chloroplast envelope and thylakoids; light green: chloroplast stroma; blue: pyrenoid Rubisco matrix; brown: starch; darker orange: eyespot granules; darker red: mitochondrial envelope and cristae; light red: mitochondrial matrix; grey: cell wall). Surrounding panels: slices through example tomograms (greyscale), with positions of the molecular complexes marked (arrowheads), and STA density maps of those complexes overlaid (in color). Scale bar: 50 nm; all tomograms shown at same scale. **A)** Nucleosomes in the nucleus. STA map: 9.6 Å resolution, purple: DNA, orange: histone H2A, red: histone H2B, blue: histone H3, green: histone H4. See also Fig. S8. **B)** Photosystem II (PSII) embedded within thylakoid membranes of the chloroplast. STA map: 19 Å resolution, green: PSII, grey: membrane. See also Fig. S11. **C)** Rubisco in the pyrenoid compartment of the chloroplast. STA map: 7.5 Å resolution, dark blue: Rubisco large subunits, light blue: Rubisco small subunits. See also Fig. S7. **D)** Microtubules in the cytoplasm. STA map: 4.7 Å resolution, light and dark green: tubulin monomers. See also Fig. S9. **E)** Clathrin lattice in the cytoplasm. STA map: 8.7 Å resolution, pink: clathrin heavy chains, blue: clathrin light chains. See also Fig. S10. **F)** ATP synthase dimers embedded in the crista membranes of mitochondria. Composite STA map: peripheral stalk (yellow) at 5.2 Å resolution, F_1_ head (shades of red) at 8.5 Å resolution, purple: F_0_ rotor ring, blue: central stalk. See also Figs. S12-S14.

#### Rubisco in the chloroplast

Rubisco enzymes perform carbon fixation, assimilating CO_2_ into sugar. Green algae such as *C. reinhardtii* contain type-I Rubisco, a ∼520 kDa holocomplex composed of eight large subunits and eight small subunits. *C. reinhardtii* increases the efficiency of CO_2_ fixation by densely packing Rubisco into a chloroplast microcompartment called the pyrenoid, which forms by liquid-liquid phase separation and is traversed by a network of membrane tubules. (Fig. 2H)^83^. The high concentration of Rubisco particles within the pyrenoid provides an opportunity to test STA workflows for relatively rigid and small particles immersed in a crowded cellular environment.

Despite the dense packing of particles, discrete Rubisco holocomplexes were clearly visible within tomograms of the pyrenoid (Fig. 3C). We implemented a template matching workflow using a template density simulated from a crystal structure^94^, which we used to pick ∼240k particles from the dataset. After STA with multireference alignment, we obtained a 7.5 Å-resolution map (Figs. 3C, S7) from a class containing ∼14k particles (6% of the initial picks). This small percentage of picked subvolumes (including many true-positive Rubisco particles from visual inspection) that made it into the final high-resolution average highlights the challenges to template matching and subtomogram alignment posed by molecular crowding *in situ*. It is also likely that the particles within the pyrenoids are more heterogeneous than we had originally expected, due to Rubisco binding partners such as Rubisco activase. Continued computational developments are required to better resolve the majority of small particles in such a crowded environment.

#### Nucleosomes in the nucleus

Like in plants and animals, the nuclear genome of *C. reinhardtii* is packaged into chromatin. The fundamental unit of chromatin is the nucleosome, an octamer of histone proteins wrapped with ∼147 base pairs of DNA^95^. Nucleosomes are exciting targets for *in situ* cryo-ET, because resolving their native structures and organization within cells has the potential to provide fundamental new insights into gene regulation. However, nucleosomes are relatively small (∼200 kDa), and the nucleus is densely packed with DNA and other macromolecules, making it difficult to unambiguously pinpoint specific structures in tomograms. Consequently, the highest resolution nucleosome structure within cells is currently limited to ∼12 Å^96–100^. *C. reinhardtii* has the advantage of a smaller nuclear genome (120 Mb compared to 6.27 Gb in human cells), and nuclear tomograms show a relatively sparse distribution of macromolecules (Fig. 3A), making it a promising model system for structural investigations of native nuclear complexes.

Using template matching and STA, we determined the nucleosome structure in *C. reinhardtii* at sub-nanometer resolution within the cell. Not only does the relatively small size of the nucleosome make it a challenging target, the wrapping DNA, which is denser than protein, causes the side views to be more prone to detection than the top and bottom views, skewing the angular distribution of the picked particles (Fig. S8G). Similar to our analysis of Rubisco, only ∼11% of particles picked by template matching were retained in our final average after classification (∼24k out of ∼224k particles), highlighting the challenge of studying small complexes within crowded environments. Nevertheless, a global resolution of 9.6 Å was achieved, revealing secondary structure elements within the histone core, including defined α-helices (Fig. S8F,H). At this resolution, the helical turns of DNA begin to be visible (Fig. 3A). We observe additional diffuse density near the nucleosome dyad and entry/exit site, known hotspots for interaction, which can likely be attributed to a mixture of bound factors. Interestingly, on one face of the nucleosome, we observe an additional α-helical density near histone H2B that cannot be explained by the existing human, yeast, or xenopus eukaryotic nucleosome models (Fig. S8I). Future studies will be required to understand the origin of this extra density, and whether it is present within native cellular chromatin organization across eukaryotes, or is specific to *C. reinhardtii*.

#### Microtubules in the cytosol

The cytoplasmic microtubules (MTs) of *C. reinhardtii* are nucleated near the centrioles and extend around the periphery of the cell. In our dataset, we observed both single MTs and bundles of 3-4 MTs, perhaps corresponding to the cortical and rootlet MTs, respectively^101,102^. MT protofilaments were clearly discernible in cross-section and side views (Figs. 3D, S9A). *C. reinhardtii* cytoplasmic MTs have 13 protofilaments, the canonical number of protofilaments that is also observed in mammalian cells^103^. This conservation is noteworthy, as a diversity of protofilament number has been observed in some other species, including 11 and 15 in *C. elegans* worms^13,104^ and 13-18 in malaria parasites^5^. Recent *in situ* cryo-ET studies of mammalian neurons and pluripotent cells have produced STA maps of 13-protofilament MTs with resolutions ranging from 12 - 8.2 Å^105,106^. We therefore proceeded to average the MTs we found in the *C. reinhardtii* dataset to benchmark the quality with which we could resolve these conserved cytoskeletal structures.

After extracting particles along traced MT filaments and performing an STA strategy that accounts for MT geometry and polarity, we obtained a 4.7 Å-resolution map of the *C. reinhardtii* MT lattice from only ∼15k subvolumes (Figs. 3D, S9B-H). The map shows clear secondary structural elements including α-helices and β-strands, as well as large aromatic side chains (Fig. S9I). Notably, our STA revealed that cytoplasmic MTs have uniform polarity orientations in all tomograms, consistent with a polarized cytoskeleton nucleated at the apical end of the cell near the centrioles. This polarity is comparable to the axonal MT cytoskeleton observed by cryo-ET of mammalian neurons^105^.

#### Clathrin coats in the cytosol

Clathrin is involved in the budding of coated vesicles on the plasma membrane, the trans-Golgi network and endosomes. Upon recruitment to the membrane by adaptor proteins, clathrin forms a lattice-like coat composed of a mixture of pentagons, hexagons, and heptagons. It has a characteristic triskelion geometry consisting of three heavy chains (∼190 kDa) and three light chains (∼25 kDa). While clathrin has not been extensively studied in *C. reinhardtii*, its sequence is well-conserved between *C. reinhardtii* and mammals, and the mammalian structure has been determined to high resolution *in vitro*^107^ and to low resolution *in situ*^108^. We identified ∼6k clathrin triskelia using template matching and subsequent cleaning. These particles were fed into an STA pipeline, yielding a final 8.7 Å-resolution map from ∼18k C1-symmetric particles (Figs. 3E, S10A-H). At this resolution, the *in situ* map is similar to the previously reported *in vitro* structures, implying high structural conservation between mammals and green algae, and confirming that the *in vitro* structures are biologically representative.

Placing the clathrin structure back into the tomograms revealed clathrin assemblies at various stages of vesicle maturation, ranging from budding clathrin-coated pits to mature clathrin-coated vesicles. These assemblies displayed a variety of clathrin lattice polygonal architectures, including pentagons, hexagons, and heptagons (Fig. S10I). In addition to visualizing and characterizing the native lattice organization, structural work on clathrin *in situ* may allow the roles of interacting factors in lattice assembly and disassembly to be assessed. The potential for localizing and resolving different components of the clathrin pathway in action within the cell remains to be explored, potentially using this *C. reinhardtii* dataset.

#### Photosystem II in the chloroplast

Resolving membrane proteins remains a major challenge for cryo-ET. The high-atomic-weight phosphate head groups of lipid bilayers create strong signal in transmission electron microscopy, which interferes with the analysis of membrane-embedded protein domains both in particle picking and the alignment of subtomograms. We therefore decided to test our *C. reinhardtii* dataset for the feasibility of resolving photosystem II (PSII), one of the most abundant membrane proteins in the chloroplast, an organelle that occupies almost half of the cell volume and accordingly constitutes a significant portion of the dataset. PSII is the starting point of the photosynthetic electron transport chain, which is responsible for converting light into biochemical energy. Functional PSII assembles into a ∼600 kDa homodimeric complex, which is found in the stacked regions of the thylakoid membranes. The bulk of PSII is membrane-embedded^109^, while the luminal domains of the complex form only a small characteristic bump visible in tomograms of thylakoid membranes (Fig. 3B)^84^, making this a challenging test target for particle picking and STA.

Our attempts with template matching failed to accurately localize PSII (Fig. S11I). Likely, strong signal arising from the membrane, as well as fringing artefacts accumulated from the repetitive architecture of stacked thylakoid membranes, interfere with obtaining clear cross-correlation peaks from template matching. To circumvent this issue, we developed a custom U-net-based picking strategy to obtain positions of 26,827 putative PSII particles from 21 high-quality chloroplast tomograms (Fig. S11A-E). We used the defined orientation of PSII relative to the membrane to determine initial angles for the particles, followed by STA to resolve this membrane protein complex to a resolution of 19 Å (Figs. 3B, SF-H). The overall shape of our STA map agrees well with PSII models obtained by single-particle cryo-EM (Fig. S11G)^109^.

#### ATP synthase in mitochondria

Mitochondria contain several large membrane protein complexes involved in cellular energy metabolism. ATP synthase is embedded in the crista membranes, where it regenerates ATP by using the proton gradient across the membrane built up by the respiratory chain. Previous cryo-ET studies have shown that ATP synthase in yeast and animal forms rows of dimers, which induce the high membrane curvature at the crista edges^110–112^. Purified F_1_F_0_-ATP synthase was previously characterized by single-particle cryo-EM from a related algal species, *Polytomella sp*^113^. Recently, this line of investigation was extended with *in situ* cryo-ET to reveal ATP synthase inside native *Polytomella* cells at sub-nanometer resolution and distinguish distinct rotary conformational states of the central stalk and the F_1_ head^114^.

To resolve the *in situ* structure of ATP synthase from *C. reinhardtii*, we performed template matching and STA of these large membrane complexes within native mitochondria (Fig. 3F, S12-S14). After determining an initial average of ATP synthase dimers (Fig. S13B), we performed consensus refinement for the peripheral stalk region, yielding a 5.2 Å map with local resolution to 4.8 Å (Fig. S13E, G). The F_1_ head was less resolved in the consensus map due to its conformational movements. We used 3D classification to discriminate different rotary states (Fig. S13C), and subsequent refinement of the most populated class yielded a map of the head region at 8.5 Å resolution (Fig. S13D, G). While canonical ATP synthase subunits (α, β, γ, δ, ε, c10, a, and OSCP) are well conserved across species beyond *Polytomella* and *Chlamydomonas*^115^, the sturdy peripheral stalk formed by ATP-synthase associated units (ASA1-10) is specific to these green algae species^116^. Dimers of ATP synthase are organized in helical arrays, with 13 nm spacing, and 12° and 16° bend and twist angles, respectively (Fig. S14A-B). Overall, the *in situ* structure and organization of ATP synthase are quite similar in *Chlamydomonas* and *Polytomella*. Both *in situ* structures contain extra density not present in the previous *in vitro Polytomella* structure^113^, which corresponds to the ASA3 subunit forming a long α-helix parallel to the membrane and ending in a spindle-shaped density contacting the a-subunit^114^ (Fig. S14D-F). However, we noted one significant difference between the *in situ* structures from these two green alga species: *Chlamydomonas* lacks the “bridge” dimer linkage between ASA1 subunits in the upper part of the peripheral stalk (Fig. 3F, S14D). Interestingly, despite this difference, the angle between the two central stalks *in situ* is nearly identical in the two species (65° in *Chlamydomonas*, 66° in *Polytomella*)^114^ (Fig. S14C).

### Data sharing and exploration for community-driven visual proteomics

EMPIAR^49^ serves as the global archive for freely accessible raw electron microscopy data of biological samples. To help drive computational development and biological discovery, we have deposited both the raw and processed data corresponding to the entire 1829 tomogram dataset into EMPIAR (28 TB in total; EMPIAR-11830). We also created a public GitHub repository with all particle positions and orientations used for the STA projects in this paper, and we encourage future depositions by the community as additional particle classes are investigated. However, it is challenging to search and visualize large cryo-ET datasets, particularly *in situ* cellular datasets that are rich in diverse biological information. To facilitate discovery, all reconstructed tomograms have also been deposited into the CZII CryoET Data Portal^50^ (DS-10302), where they can be individually browsed online in a 3D volume viewer, along with annotations for particle positions and membrane surfaces (the latter generated automatically with MemBrain software)^45^. We invite scientists, educators, and students to interactively explore these tomographic volumes and accompanying annotations of the native cellular environment.

## Discussion

Our study demonstrates the application of modern cryo-PFIB and cryo-ET instrumentation to produce a large-scale dataset of high-quality tomograms (Fig. 1), which provide a detailed view of the molecular architecture inside native *C. reinhardtii* cells (Fig. 2). The complexity of biology captured in these cellular tomograms highlights the immense potential of *in situ* cryo-ET, but also the significant challenges to understanding this complexity that can only be overcome through collective community efforts. We therefore provide this dataset as an open community resource, with the hope that it will enable new biological discoveries and the continued development of computational methods.

There are many biomolecular complexes awaiting structural study in this *C. reinhardtii* dataset. To verify the utility of our dataset for this purpose, we performed STA trials of seven abundant complexes (80S ribosome, Rubisco, nucleosome, microtubule, clathrin, PSII, ATP synthase), which varied in size (>3 MDa to ∼200 kD) and cellular localization (Figs. 1 and 3). Six of these complexes reached sub-nanometer resolution, underscoring the exciting potential of modern cryo-ET workflows to resolve a variety of structures within their native cellular context. Each average was generated by a different user, each with their own STA pipeline using a distinct combination of software (see methods).

These STA trials highlight challenges that require further exploration and computational developments. In most cases, only ∼20k particles were required to reach sub-nanometer resolution. However, in the case of small particles such as Rubisco and the nucleosome, this required classification steps that discarded 89-94% of the particles originally selected by template matching. Template matching algorithms are less effective for detecting small particles, which provide less signal, and downstream classification routines can struggle to cleanly sort the extracted subvolumes into different class averages. This high discard rate may be partially due to biological heterogeneity (especially in the case of the nucleosomes), but also reflects the inherent properties of noisy anisotropic cryo-ET data, which make it difficult to identify all true positive instances of a target macromolecular species without also selecting false positives. These issues with template matching and classification are problematic if one wants to capture an accurate census of the macromolecules within the cellular volume, a central goal of visual proteomics. For example, while it is feasible to generate high-resolution averages of nucleosomes and Rubisco complexes through extensive classification, mapping only this small subset of true positive particles back into the cellular volumes cannot provide accurate information about the molecular organization of chromatin or the pyrenoid matrix.

Further complicating these technical challenges is the biological challenge of the crowded native cellular environment, itself. Cells present landscapes of densely packed, and even directly contacting, particles in a myriad of conformational states. Promisingly, cryo-ET method development is very active in the areas of feature detection^44,46,117–120^, particle picking^47,121–123^, and heterogeneity analysis^64,124–126^, with many new approaches leveraging neural networks. These AI-based programs benefit from large quantities of training data, which our *C. reinhardtii* dataset can help provide. Membrane proteins can be especially challenging *in situ*, exemplified by our subtomogram average of PSII. Only a relatively small portion of the membrane complex extends from the density of the membrane, and the stacked thylakoid membranes themselves are exceptionally straight, resulting in strong CTF effects that complicate particle detection and alignment. Specialized computational workflows are under development for the analysis of membranes and their embedded proteins^45,127^, including the MemBrain program, which benefited significantly from the *C. reinhardtii* dataset to improve the training of membrane segmentation.

To realize the goals of visual proteomics, cryo-ET analysis must dig deeper to resolve the rich diversity of molecular complexes inside cells. Especially for less abundant complexes, this goal would strongly benefit from large repositories of publicly available high-quality data that are easy to search, explore, and combine. *In situ* cryo-ET datasets are dense with information that has uses well beyond the specific aims of a single study or research group; the challenge is to establish best practices and new tools that will unlock the value of all that unused visual proteomics information. Continued efforts to facilitate the process of data sharing within the cryo-ET community can help bring about the next generation of AI-driven computational methods. Combined with parallel advancements in mass spectrometry and structure prediction^2^, these methods may be able to provide deep inventories of diverse biomolecular complexes in tomograms, revealing their structures, their conformational heterogeneity, their interactions, and their 3D molecular organization inside native cells.

## Material and Methods

### Sample Preparation

Cells used in this study were *Chlamydomonas reinhardtii* strain mat3-4 (CC3994), acquired from the Chlamydomonas Resource Center, University of Minnesota, MN. This strain has a smaller cell phenotype^128^, which is advantageous for vitrification. From an agar plate stock culture, a liquid culture was inoculated in 40 mL Tris-acetate phosphate (TAP) media, containing 0.02 M Tris base, TAP salts (7.00 mM NH_4_Cl, 0.83 mM, MgSO_4_, 0.45 mM CaCl_2_), Phosphate buffer (1.65 mM K_2_HPO_4_, 1.05 mM KH_2_PO_4_), and Hunter trace elements (0.134 mM Na_2_EDTA, 0.136 mM ZnSO_4_, 0.184 mM H_3_BO_4_, 40 µM MnCl_2_, 32.9 µM FeSO_4_, 12.3 µM CoCl_2_, 10.0 µM CuSO_4_, 4.44 µM (NH_4_)_6_MoO_3_, 17.5 mM acetic acid). Cells were constantly illuminated (∼90 µmol photons/m^2^s), agitated at 100 rpm in normal atmosphere, and harvested 48–72 h after inoculation. A total of 4 µL of cell suspension (1500–2000 cells/µL) were applied to holey carbon R2/1 or R1/4 copper TEM grids (Quantifoil), and cells were plunge frozen on a Vitrobot Mark IV (Thermo Fisher Scientific, settings: blot force = 10, blot time = 10 s, temperature = 30 °C, humidity = 90%) using back-side blotting and either an ethane/propane or an ethane bath. Samples were stored in liquid nitrogen until use.

### Serial cryo-FIB/SEM volume imaging

Sample grids were clipped in Autogrids (Thermo Fisher Scientific) and subjected to automated serial slice FIB/SEM volume imaging using Auto Slice & View 5 (Thermo Fisher Scientific) in a Hydra Bio PFIB dual beam instrument (Thermo Fisher Scientific). Grids were loaded using the standard 35° pre-tilted holder, then coated with conductive and protective layers including: Pt sputter (magnetron, 30 mA, 10 pA, 30 sec); organometallic Pt condensate using gas injection system (2 min); Pt sputter (magnetron, 30 mA, 10 pA, 30 sec). For serial FIB/SEM volume imaging, these samples were tilted such that the grid is flat and perpendicular to the electron beam (i.e., 35° stage tilt), resulting in a FIB milling angle of 38°. Sites were milled using argon with 30 kV accelerating voltage and 200 pA nominal beam current. Slice thickness was 20 nm. SEM imaging conditions were optimized with 1.2 kV accelerating voltage and 13 pA beam current using immersion mode with TLD-DH detector settings. Image collection parameters were 50 ns dwell time with 150 line integrations and a 5 x 5 nm pixel size.

### Cryo-FIB lamella preparation

Grids were clipped into Autogrids (Thermo Fisher Scientific) and subjected to automated lamella preparation using an Arctis cryo-plasma-FIB (Thermo Fisher Scientific) with two different workflows. For the first workflow, prior to milling, grids were coated with a layer of ion-sputtered, metallic platinum (Pt) for 120 sec (Xe+, 12 kV, 70 nA). This was followed by 400 nm organometallic Pt using the gas injection system, then an additional ion-sputtered platinum layer (Xe+, 12 kV, 70 nA, 120 sec). Next, grids were surveyed using Maps software (Thermo Fisher Scientific) for lamella site identification followed by automated lamella preparation using AutoTEM Cryo with a thickness range set between 150 – 300 nm for polishing steps. The second workflow was performed with Arctis WebUI (Thermo Fisher Scientific). Using the WebUI interface, grids were screened automatically, coated with the same amount of Pt sputtering-GIS-sputtering as in the first workflow, then subjected to automated lamella preparation. All FIB milling was performed using xenon or argon ions.

### Cryo-ET data acquisition

Datasets were collected using a Krios G4 equipped with a Selectris X energy filter and Falcon 4i direct electron detector (Thermo Fisher Scientific). Tilt-series were collected with a dose-symmetric tilt scheme using TEM Tomography 5 software (Thermo Fisher Scientific). The tilt span of ± 60° was used with 2° or 3° steps starting at either ± 10° to compensate for the lamella milling angle. Target focus was set for each tilt-series in steps of 0.25 µm over a range of -1.5 µm to -3.5 µm. Data was acquired in EER mode of Falcon 4i with a nominal pixel size of 1.96 Å and a total dose of 3.5 e^−^/Å^2^ per tilt. A 10 eV slit was used for the entire data collection. Eucentric height estimation was performed once for each lamella using stage tilt method in TEM Tomography 5 software. Regions of interest were added manually, and positions saved. Tracking and focusing was performed before and after acquisition of each tilt step. The energy filter zero-loss peak was tuned once before each data acquisition session.

### Preprocessing

The data were preprocessed using TOMOgram MANager (TOMOMAN)^129^. EER images were motion corrected using a modified version (to generate odd-even frame sums) of RELION’s implementation of MotionCor2^130^. The defocus was estimated using tilt-corrected periodogram averaging in tiltCTF^131^ along with CTFFIND4^132^ implementation within TOMOMAN. Tilt-series were aligned using fiducial-less alignment in AreTomo^133^. Initially, tomograms without CTF correction were reconstructed by IMOD’s tilt program. Subsequent odd-even reconstructions and denoised tomograms were generated using cryo-CARE. 3D-CTF corrected reconstructions were generated with novaCTF using phase-flipping algorithm in batch. All tomographic reconstructions were downsampled (binned) using the Fourier3D program (https://github.com/turonova/Fourier3D) when applicable. Exporting to Warp-RELION-M^32,134^ and RELION-4^135^ tomography workflows was performed using TOMOMAN for STA. All operations were performed in batch mode as implemented in TOMOMAN.

### Lamella thickness measurement

Lamella thickness was measured in each reconstructed tomogram (Fig. 1E) through a TOMOMAN script based on IMOD’s findsection program^136^. In cases where findsection failed to estimate a boundary model due to low tomogram contrast, an indirect measurement was made based on boundary masks automatically generated with Slabify^137^.

### Regression Denoising

Synthetic cryo-ET data for regression U-net training was generated using CryoTomoSim^88^. Using a box of 400 x 400 x 50 pixels, a mixture of medium and small proteins were modelled in four layers at 7.84 Å/px. The exact protein identities are not important to training. Ten membrane vesicles were modelled. Particle density was 0.8, the vitreous ice option was used, and the iterations for each layer were 500, 500, 4000, 8000. The output (“ideal” tomogram) was simulated at -1 µm defocus, -89 to 89 tilt, 0.5 degree tilt increment and total dose = 0. The input (noisy tomogram) was simulated at -3 µm defocus, -60 to 60 tilt, 3 degree tilt increment, and total dose = 80. Both datasets were loaded into Dragonfly 2022.2 (Object Research Systems) and used as training input and output for a regression 2.5D U-net with the following architecture: depth level 5, initial filter count 64, slice count 5, patch size 128, Loss function ORSMixedGradientLoss. Training proceeded for 46 epochs using a total of 15,840 patches.

### Segmentation

A combination of shallow and 2D U-nets were trained using Amira 2023.2 (Thermo Fisher Scientific) to segment ribosomes, COPI, nuclear pore complexes, microtubules (singlets and triplets), basal body cartwheel, and Rubisco in Fig. 2. Approximately 200 patches from each tomogram were selected after hand annotation for AI-assisted segmentation. The resulting segmentation was used for further patch extraction to train a U-net of the following architecture: 2D, backbone: resnet18, patch size: 128, loss function: dice. All membrane segmentations for display (Figs. 2C-H, S6A-D) were obtained using the MemBrain-seg module of the MemBrain v2 package^45^. ATP synthase (yellow label in Fig. 2F, blue label in Fig. S6A-B), nucleosomes (pink label in Fig. 2C), and ribosomes (orange label in Fig. S6A-B) are particle mapbacks of the STA maps at template matching coordinates. The 2.5D segmentation U-net for PSII was trained using Dragonfly 2022.2 (Object Research Systems) according to a previously described protocol^89^. In brief, Segmentation Wizard was used for manual annotation of 5-6 training slices, and a generic U-net of the following architecture was trained: 2.5D: 3 slices, depth level: 5, initial filter count: 64, patch size: 128. Training data included slices from tomograms numbered 24, 373, 473, and 900. Aside from patch size, all hyperparameters were left as default. Training labels were membrane, ATP synthase, ribosomes, PSII and background. The 2.5D segmentation U-nets for Fig. S6 segmentations were trained using Dragonfly 2022.2 (Object Research Systems) according to the same protocol using the following architecture: Pretrained U-net, 3-slices, depth level: 5, initial filter count: 64, patch size: 96. Unique U-nets were trained for each segmentation.

### STA: 80S Ribosome

Initial ribosome positions and orientations were determined by template matching using STOPGAP^29^ or its GPU implementation GAPSTOP-TM^27^ (https://gitlab.mpcdf.mpg.de/bturo/gapstop_tm) on ∼1000 8x binned tomograms, resulting in ∼500k particles. These were then iteratively aligned using STOPGAP at 8x and 4x binning using a soft mask shaped to contours of the full ribosome density. Particle scores were distributed bimodally; the ∼140K particles in the higher-scoring distribution were selected for further processing. These particles were further aligned at 2x binning, first using a full ribosome mask, followed by alignment using a mask focused on the large subunit. The resulting orientations were used as a starting point for two separate multireference alignment (MRA) runs in STOPGAP (Fig. S2E-G). Starting references were generated *de novo* by randomly assigning 20% of the dataset to ten classes. To classify the membrane bound state, MRA was performed with a mask focusing on the membrane bound region. To classify the different tRNA states, MRA was performed with a mask focusing on the tRNA channel. In both cases, the first 10 iterations of MRA were performed using simulated annealing, followed by MRA with stochastic hill climbing and without simulated annealing until convergence, i.e., when less than 1% of subtomograms changed classes between iterations. Final classes were assigned by visually inspecting the class averages and merging identical states. This resulted in five unique states: ‘A, P unrotated’, ‘A, P rotated’, ‘eEF2, A/P, P/E rotated’, ‘eEF2, P, E partial rotated’, and ‘eEF1A, A/T, P unrotated’^138^ (Fig. S2G). For higher resolution averaging, particles were exported to Warptools^139^ at 1x binning (1.96 Å/px) using TOMOMAN^129^. After importing, three iterations of M were performed with 2D image warp refinement, particle pose refinement, and exhaustive CTF refinement. Of these, the last iteration was refined for tilt angles. This resulted in the 80S ribosome density map at global resolution of 4.0 Å from ∼140K particles from 600 tomograms (Figs. 1F, S2D). For analysing the relationship between Rosenthal-Henderson B-factor and lamella thickness (Figs. 1G, S3), particles were segregated based on the thickness of the corresponding tomogram. Thickness bins were defined as 100-125, 125-150, 150-175, 175-200, 200-225, 225-250, >250 nm. To analyse the damage due to FIB milling (Figs. 1H, S4), we calculated Rosenthal-Henderson B-factor for particles based on the distance of the particle centre (depth) from the edge of the lamella (tomogram boundary model). Depth bins were defined as 0-15, 15-30, 30-45, 45-60, 60-75, 75-90, 90-105, >105 nm from the lamella surface. In both these cases, we used the boundary models created during the tomogram thickness measurements (see “lamella thickness measurement” section above). For each thickness and depth bin, Rosenthal-Henderson B-factor was calculated based on the linear fit of resolution^−2^ vs. ln(number of particles) for subsets with 2000, 4000, 6000, 8000, and 10000 particles (Figs. S3, S4).

### STA: Rubisco

The template map was generated from PDB: 1gk8^94^ using the “simulate” program within the cisTEM package^140^. This map was used to determine initial Rubisco positions and orientations by template matching in STOPGAP on 16 4x binned tomograms (Fig. S7A-B), resulting in approximately ∼240k particles. These subvolumes were then iteratively aligned using STOPGAP at 4x and 2x binning using a soft mask shaped to contours of the full Rubisco holocomplex density. The resulting orientations were used as a starting point for an MRA protocol in STOPGAP. After classification, we obtained a 7.5 Å class average from ∼14k particles (Figs. 3C, S7C-D).

### STA: Nucleosomes

126 tomograms from the curated dataset containing nuclei in the field of view were selected for processing based on their visual quality. For particle picking, first with a subset of 8 tomograms, we applied STOPGAP^29^ for template matching on 4x binned tomograms. The template map was generated from PDB: 3AFA^141^ at 7.84 Å/px using the molmap command in ChimeraX^142^. Subtomograms were reconstructed at 3.92 Å/px (2x binning) in Warp and then subjected to iterative rounds of alignment and classification, followed by 3D auto-refinement in RELION v3.1.4^143^. The resulting nucleosome map (∼15 Å) was then used as a template to pick particles within the entire set of 126 tomograms using the GPU-accelerated version of PyTom^30^ for template matching (https://github.com/SBC-Utrecht/pytom-match-pick) on 4x binned tomograms (Fig. S8A-B). In order to exclude obviously false cross-correlation peaks outside the nucleus, nuclear masks were predicted by a neural network using MemBrain-seg^45^, which was trained with 8x binned segmentations manually created with Amira (Thermo Fisher Scientific). Up to 8,000 putative particles were extracted from each tomogram following standard PyTom template matching procedures, yielding a total of 223,507 particles. Subtomograms were reconstructed at 3.92 Å/px (2x binning) in Warp and then subjected to multiple rounds of 3D classification and 3D auto-refinement in RELION v3.1.4^143^. The best class, containing 23,878 particles (Fig. S8C-D), was then exported back to Warp^134^ for subtomogram extraction using custom TOMOMAN scripts^129^. Subsequently, multi-particle refinement was performed in M^32^ to refine particle poses as well as parameters of the tilt-series geometry and the contrast transfer function (CTF). Refinements in M and RELION were iterated at 2x and 1x binning (1.96 Å/px) until convergence, yielding a nucleosome map at a global resolution of 9.6 Å according to the Fourier shell correlation 0.143 threshold^144^ (Figs. 3A, S8E-I). The final refinement in M was performed with an input mask around the histone octamer core, excluding the DNA.

### STA: Microtubules

The STA procedure for the MTs was performed essentially as described before^105^. MTs in 8x binned tomograms (15.68 Å/px) were segmented with the automated template-matching procedure in Amira software v.6.2.0 (Thermo Fisher Scientific) as previously described^145,146^ using a generic template of a missing wedge-modulated 120 nm long hollow cylinder with outer and inner radii of 14 and 7 nm, respectively (Fig. S9A-B). The coordinates of the traced MT center lines were further resampled in MATLAB (MathWorks) to obtain equidistant points at every 3.2 nm. Resampled coordinates were used to generate a tubular grid around the center line and place points approximately on each tubulin subunit taking into account inter-subunit distance of 40 Å along the filament z-axis and 60 Å around the helix. These positions were oversampled 2x, and subtomograms were extracted from these grid points. Each MT in a given tomogram was assigned a class number, which helped in determination of MT polarity. Subtomogram averaging was performed in STOPGAP^29^. An initial reference was first generated de novo from a tomogram containing MTs that have the same polarity orientations. 10,276 subtomograms were extracted from the oversampled positions using 2x binning (7.84 Å/pixel) and a box size of 64 px³. A starting reference was generated by averaging all subtomograms. Several rounds of alignment of the in-plane angle were conducted with shift refinements limited to 4 and 6 nm in x- and y-direction along the filament center line. This averaging approach resulted in a satisfactory MT reference. We used this optimised starting model for refining the whole dataset comprising 47 MTs from 18 tomograms sequentially at 4x binning (6 iterations), 2x binning (5 iterations), and 1x binning (5 iterations). MT polarity orientations were determined by averaging each MTs separately using multiclass averaging in STOPGAP, and subtomogram orientations were visualised in Chimera^147^ as lattice maps using the Place Object plugin^148^. Polarities were sorted out using visual inspection of the tubulin radial skew^149^, then all MTs were unified to the same polarity by 180° change of the in-plane angles in the motive list. In a few occasions, radial skew could not be determined unambiguously (Fig. S9D), and therefore, these MTs were discarded. Finally, 27,251 unbinned subtomograms (1x binning, 1.96 Å/pixel) were extracted using a box size of 96 pixels³ and aligned iteratively, resulting in a refined map of a MT at a global resolution of ∼9 Å (Fig. S9E). The coordinates were then imported into Warp, extracted at 4x binned pixel size and subjected to 3D refinement in RELION-4^135^, followed by refinement in M^32^. In total, 4 rounds of refinement were performed, gradually adding parameters: tilt-series/tomogram 2D/3D geometry, particle poses, stage angles, CTF grid search and defocus. Each M refinement round was cycled with a 3D refinement in RELION-4. In the final two rounds of M refinement, dose fractions was refined in addition to previous parameters. Finally, the last 3D refinement using 1.25x binned subtomograms was performed in RELION-4, yielding an STA map at 4.7 Å overall resolution (Figs. 3D, S9G-I).

### STA: Clathrin

Initial clathrin positions and orientations were determined using template matching with the PyTom package^30^ (Fig. S10A-B). The angular search was conducted with a 3-degree step against 94 4x binned tomograms. One of the published EM maps of a clathrin hub (EMD-0122)^107^, low pass filtered to 15.7Å (Nyquist at 4x binning) in an 80-pixel box (7.84 Å/px), was used as a template. To minimise the number of false positives, membranes were masked out using MemBrain-seg membrane segmentations^45^. Remaining false positive picks were manually removed using the ArtiaX package^150^ for ChimeraX^151^. Picks that did not form a characteristic clathrin lattice (Fig. S10C) were considered false positives. This yielded 5999 clathrin triskelia. Aligned tilt-series and particle coordinates were transferred to Warp (v1.0.9)^134^, where CTF estimation was performed, and subtomograms, along with the corresponding per-particle CTF models, were extracted at 3x binning (5.9 Å/px). Subsequent refinement of clathrin triskelion was carried out in RELION (v3.0) producing a 20 Å-resolution initial STA map (Fig. S10D). The refined coordinates and orientations of the particles were used to refine the tilt-series alignment in M (v1.0.9)^32^. New subtomograms were extracted at 2x binning (3.92 Å/px), and particle positions were refined in RELION resulting in a 12 Å map. These particles were transferred back to M for another round of refinement of the tilt-series alignment. New subtomograms were then extracted at 1x binning (1.96 Å pixel size), and particle positions were refined in RELION, yielding a 9.5 Å map (Fig. S10E). The particles were once more transferred to M for symmetry expansion (C3 to C1, yielding 17,997 particles) and multispecies co-refinement with 80S ribosomes, resulting in the final STA map of clathrin at 8.7 Å resolution (Figs. 3E, S10F-H). Refining selected classes from 3D classification did not achieve a higher resolution, hence all particles were used for the final reconstruction.

### STA: Photosystem II

Particles were first picked by template matching using pytom-match-pick^30^ on 4x binned CTF phase-flipped tomograms, using 7 degree rotational search with a template map generated from PDB: 6KAC, low-pass filtered to 15.68 Å (Nyquist at 4x binning). Visual inspection revealed that this template matching procedure was unable to pick some obvious PSII particles (Fig. S11I), likely due to the relatively small soluble portion of PSII protruding out of the thylakoid membranes. Subsequently, a generalized 2.5D U-net was trained to segment PSII in four 4x binned tomograms using Dragonfly^89^ (Fig. S11A-D). Initial X, Y, Z coordinates of PSII positions were extracted as the center-of-mass from separate segmented instances of the U-net binary map, resulting in 26,827 initial coordinates. Thylakoid membranes were then segmented from the tomograms using MemBrain-seg^45^. The normal angle of each particle with respect to the membrane surface was assigned using a custom Python script, and particle positions were projected onto the membrane to obtain an initial alignment and angle assignment (Fig. S11E). Particles picked at a distance of more than 78.4 nm from the membrane were discarded. Based on these coordinates and assigned angles, 22,895 subtomograms were extracted as 4x binned subtomograms in STOPGAP^29^. Particles were classified into 4 classes (Fig. S11F), allowing for alignment with full in-plane rotational sampling, using an alignment mask focused on the luminal densities of PSII. Positions and angles from the class corresponding to PSII (13.6%, 3122 particles) were then ported into RELION-4^135^. The structure was refined with C2 symmetry and a shaped mask focused on the PSII core proteins. Per particle CTF refinement and tomogram frame alignment were performed on 1x binned subtomograms. The refined structure was filtered to the final 19 Å resolution, as determined by the 0.143 FSC threshold from independently refined half-maps^152^ (Figs. 3B, S11G-H).

### STA: ATP synthase and multispecies refinement with 80S ribosome

Nine tomograms reconstructed using IMOD, as elaborated in the pre-processing section, were used for initial processing. Template matching was performed in STOPGAP using 8x binned (80S ribosome) and 4x binned (ATP synthase) tomograms. Initial templates were generated by the simulate function of the cisTEM package, using pdb models: 6gqv (80S ribosome)^153^ and 6rd4 (ATP synthase)^113^. The template matching settings were as follows: lowpass filter to 31.4 Å and 15.7 Å (the respective Nyquist frequencies) and angle scanning of 15° and 12°, respectively. For the 80S ribosome, subtomogram averaging in STOPGAP was performed using 8x binned subtomograms to obtain a reference used for template matching of the whole dataset. For ATP synthase, 3D classification without alignments was performed in STOPGAP using 4x binned subtomograms. Multiple classes for ATP synthase as well as membrane mis-picks were used for the subsequent template matching scheme on the whole dataset to suppress false positives (Fig. S12). Next, 261 tomograms of mitochondria were selected for further processing. Raw EER frames were converted to TIFF stack containing 5 dose fractions using RELION-4. Motion correction and CTF estimation was performed on the TIFF stacks in Warp 1.0.9^134^. Tilt-series metadata in mdoc format and tilt-series alignments from AreTomo were imported into Warp to reconstruct tomograms at 8x and 4x binning. Template matching was performed using the GPU-accelerated implementation of PyTom^30^ using 8x binned (80S ribosome) and 4x binned (ATP synthase) tomograms, with the following settings: lowpass filter 31.4 Å and 15.7 Å (the respective Nyquist frequencies), and angle scanning of 11° and 7°, respectively. For ATP synthase, a scoring function to remove the background from membrane hits was generated in Matlab R2022b using the score map obtained for the ATP synthase class (Fig. S12E, class II) and correction for false positives by score maps obtained membrane-containing classes (Fig. S12E, classes III & IV). Details of the scoring function, including an example, are shown in Fig. S12. Using the positions determined by template matching, subtomograms were extracted in Warp at 4x binning in both cases and used for 3D refinement in RELION-4. Subsequently, the respective species for multiparticle refinement were created in M^32^. Masks were prepared by binarization of shaped masks created in RELION. In total 5 rounds of Multispecies co-refinement of ATP synthase and 80S ribosomes were performed, gradually adding up refinement parameters. In the initial round, only tilt-series/tomogram 2D/3D geometry was optimized. Up to this point, C2 pseudo-symmetry was used in all processing steps with ATP synthase. Next, subtomograms of ATP synthase were extracted at 1.6x binning, the particle list was symmetry-expanded (C2 to C1; doubling the number of particles), and 3D refinement was performed in RELION-4^135^ using a mask focused on the peripheral stalk. 3D classification was then performed in RELION-4 to exclude lower-quality particles, followed by another round of 3D refinement. ATP synthase species was then re-imported for a second round of multispecies co-refinement in M, CTF parameters were refined in addition to geometry using a tight binary mask around the peripheral stalk for ATP synthase. Next, subtomograms were extracted at 1.25x binning and 3D refinement was performed in RELION-4. The refined ATP synthase species was reimported to M, and masking of 80S was changed from the whole ribosome to only the large subunit. Using the updated species, two rounds of co-refinement were used to optimize the motion correction in TIFF dose fractions in addition to previously refined parameters. In the next iterative cycle, 1.25x binned ATP synthase subtomograms were extracted, followed by 3D refinement in RELION-4. At this point, we performed 3D classification masking the head domain in an attempt to resolve rotational states (Fig. S13C). One class after further 3D refinement yielded a map at 8.6 Å (∼11,000 particles), where densities in the head domain were resolved (Fig. S13D). In the final refinement in M, we used all of the previous options except refinement of motion in dose fractions. Then, particle coordinates were re-centered at a single ATP synthase monomer and unbinned subtomograms were extracted from M. Finally, 3D classification and 3D refinement were performed in RELION-4, yielding a structure of the peripheral stalk at 5.2 Å (∼88,000 particles) (Fig. S13E). 80S ribosome particles were subjected to 3D refinement in RELION-4 only once and all consecutive refinements were performed in M during the 5 rounds of multispecies refinement with ATP synthase.

### Data structure of the deposition on EMPIAR

The 28 TB of EMPIAR-11830 have been deposited with the following data structure (see Fig. S15). The data are partitioned into two image sets, one containing a minimal TOMOMAN project^129^ and the other containing 4x binned denoised tomograms for visualisation. Each tilt-series is kept in a separate subfolder named with date of data collection, location, and microscope type. Further, each tilt-series folder is split into subfolders which include raw tilt-series and auxiliary files to provide a rapid overview. The reconstructed tomograms are numbered and contained in one single folder, and a STAR file^154^ is provided that indicates numbers corresponding to the tilt-series names. We maintained the accompanying metadata according to the structure commonly used for cryo-EM datasets.

## Data availability

Raw electron microscopy data are available at the Electron Microscopy Public Image Archive (EMPIAR) under accession code EMPIAR-11830. Annotation and processing information for all 1829 tomograms is provided in spreadsheet format^155^. The following subtomogram averages have been deposited at the Electron Microscopy Data Bank (EMDB): 80S ribosome (EMD-51847), nucleosome (EMD-19906), photosystem II (EMD-51731), Rubisco (EMD-51848), microtubules (EMD-51804), clathrin (EMD-51789) and ATP synthase (EMD-51802). Particle positions and orientations used for STA are available on GitHub (https://github.com/Chromatin-Structure-Rhythms-Lab/ChlamyAnnotations). Reconstructed tomograms and annotations are also available to explore interactively at the CZII CryoET Data Portal (DS-10302, https://cryoetdataportal.czscience.com/datasets/10302/). Raw data for cryo-PFIB/SEM slice-and-view of a whole *C. reinhartii* cell has also been deposited (EMPIAR-11275).

## Author contributions

Cell growth and vitrification by W. Wietrzynski, S.K., F.W., C.L.M, & S.v.D; FIB milling by R.K., X.Z., & A.K.; tilt-series acquisition by X.Z., M.O, & A.K.; data preprocessing and reconstruction by S.K., & R.D.R., with support from W. Wan; data cleaning, curation, and annotation by S.K., R.D.R., A.K.M., S.v.D., F.W., C.L.M., P.V.d.S., H.v.d.H., & W. Wietrzynski; volume segmentation by J.H., M.O., L.L, & S.Z.; subtomogram averaging by S.K. (ribosome and Rubisco), M.O., S.K., & F.W. (ATP synthase), S.C. & M.O. (microtubule), G.T. (clathrin), A.K.M., M.O., & R.D.R (nucleosome), S.v.D, R.D.R., J.H., & L.L. (PSII); curation of particle positions on GitHub by P.H.; figure composition by S.K., R.D.R., J.H., M.O., S.C., S.K., G.T., A.K.M., S.v.D., A.K., & B.E.; manuscript writing and editing led by B.E. & A.K., with support from J.H. and contributions from all authors; supervision by J.A.G.B, J.M.P., B.D.E., & A.K.

## Acknowledgments

Calculations were performed at the Max Planck Institute of Biochemistry and the Raven Supercomputer of the Max Planck Computing and Data Facility (MPCDF) in Garching, Germany, at the sciCORE (http://scicore.unibas.ch/) scientific computing center at the University of Basel, Switzerland, and at Thermo Fisher Scientific, in Eindhoven, the Netherlands.

---

**Supplementary Figure 1.**
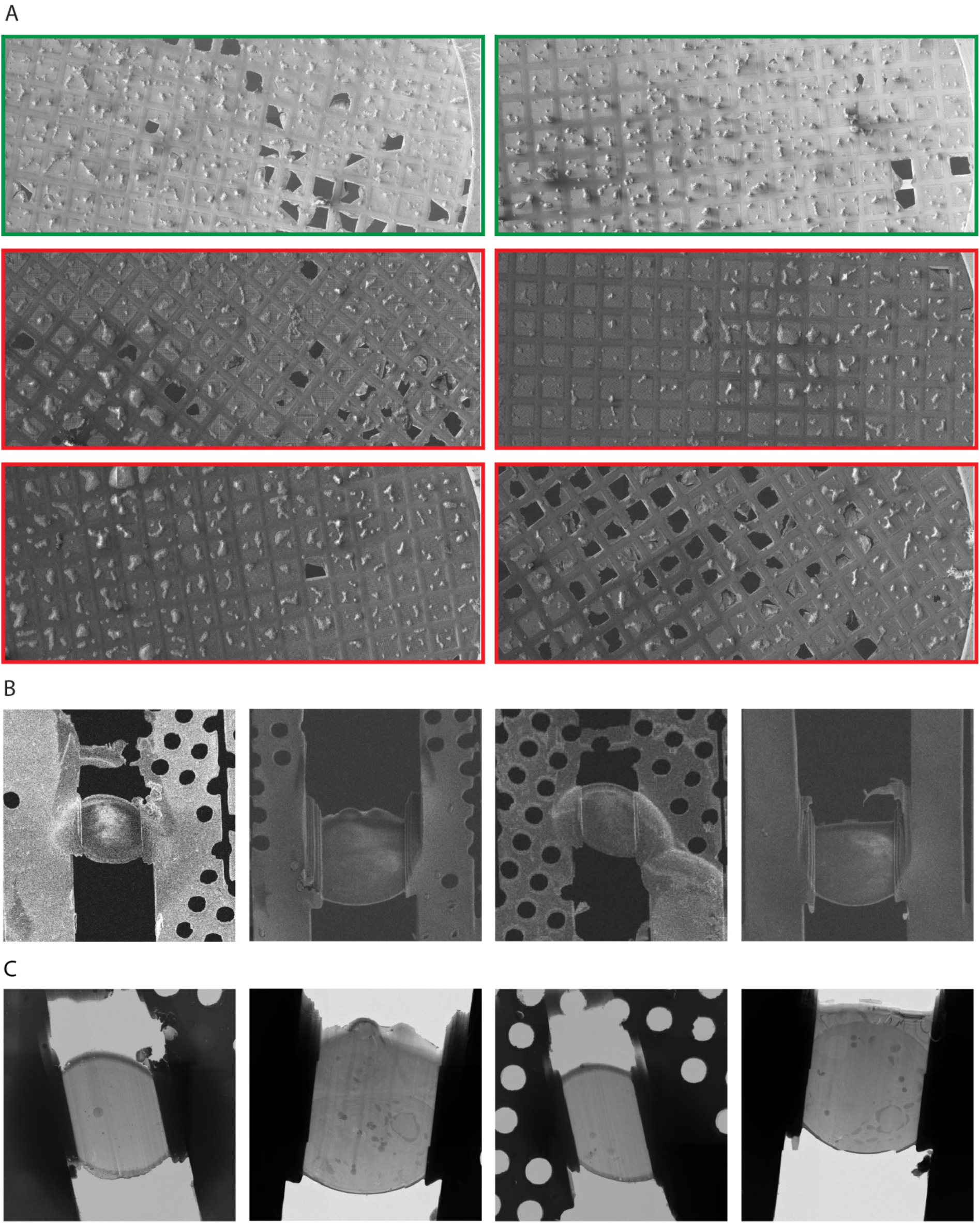
SEM overview gallery and lamella orientation in SEM and TEM. **A)** Tileset of individual grids screened with an autoloader system within the Arctis Cryo-Plasma-FIB. Green highlighted grids are high quality and were kept for milling; red highlighted grids were discarded due to poor cell distribution. **B.** SEM overviews of lamellae after final milling. **C.** TEM overviews of the same lamellae. Related to Fig. 1A-D.

**Supplementary Figure 2.**
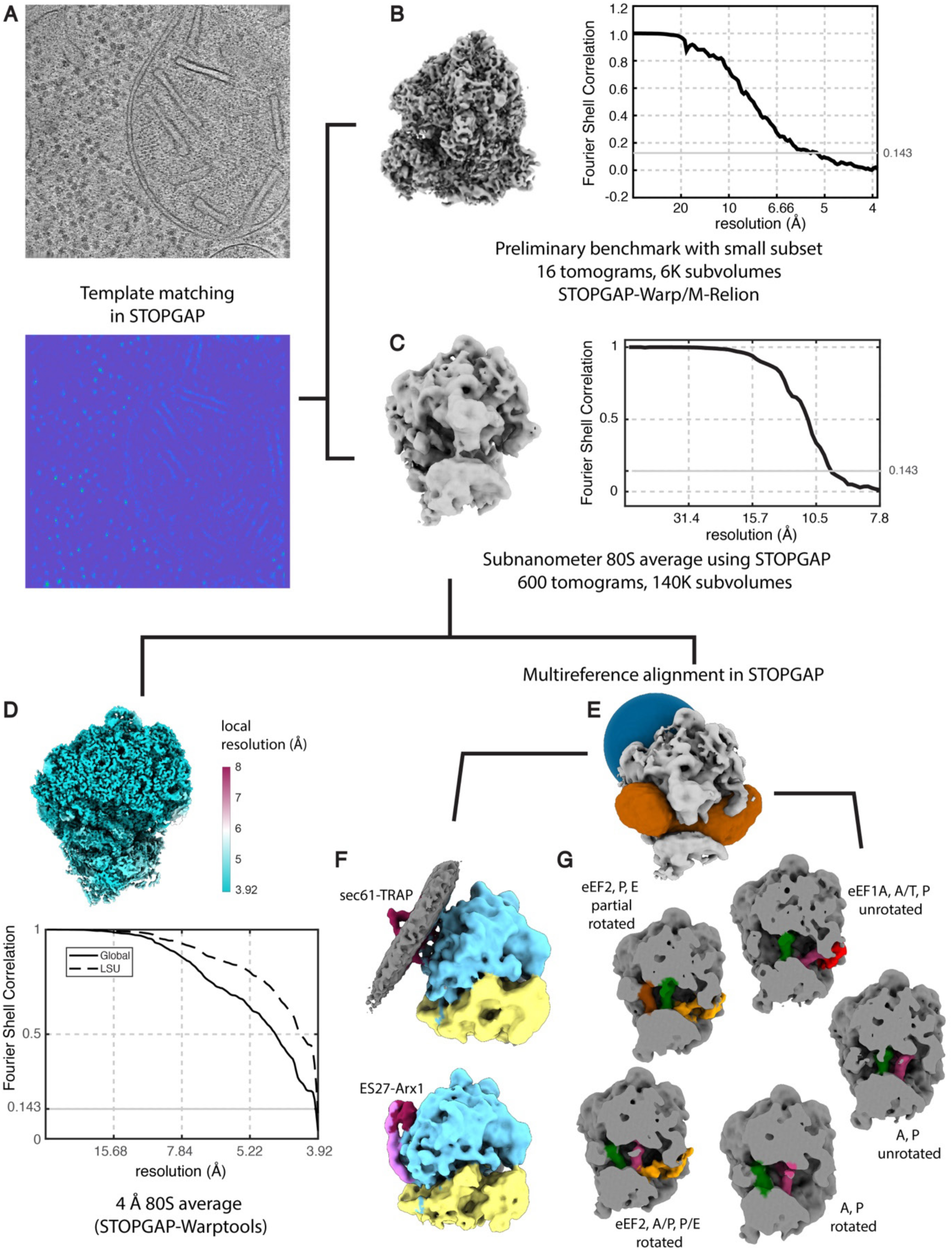
Supporting information for *in situ* STA of 80S ribosomes. Order of the processing workflow is illustrated with lines. **A)** Slice through a tomogram (top) and corresponding STOPGAP template matching score map. Green and yellow pixels: higher cross-correlation scores, blue: low cross correlation. **B)** STA map and Fourier shell correlation (FSC) global resolution estimation (5.5 Å resolution), using a test set of ∼6,000 subvolumes and the full STOPGAP -Warp/M - RELION processing workflow. **C)** Initial STA map and FSC plot (9.7 Å resolution) using a large set of ∼140,000 subvolumes averaged in STOPGAP. **D)** Final STA map from Fig. 1F (shown sliced into the interior of the structure with local resolution extending to Nyquist) and global FSC (4.0 Å resolution). **E)** Local masks used to guide multireference alignment in STOPGAP. Blue mask yielded the averages in panel F, orange mask yielded the averages in panel G. **F)** Classification of ER-bound ribosomes (blue: large subunit, yellow: small subunit, red: sec61-TRAP, grey: membrane) from free cytosolic ribosomes (magenta: ES27, red: Arx1) **G)** Classification of ribosomes translational states. Red: eEF1, orange: eEF2, pink / green / brown: tRNA moving through the A, P, and E positions.

**Supplementary Figure 3.**
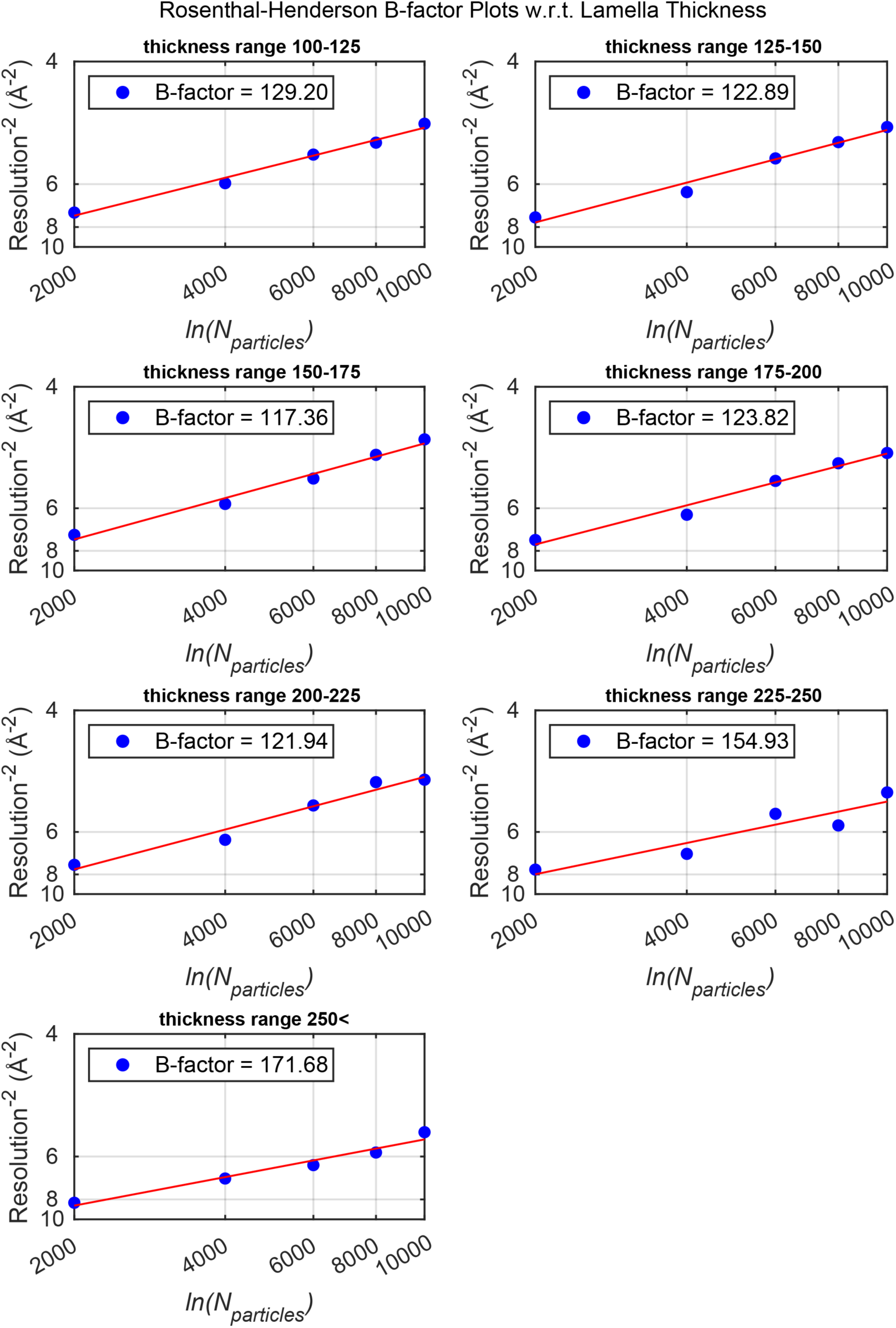
Rosenthal-Henderson B-factor plots with respect to lamella thickness. B-factor is calculated based on the linear fit of resolution^−2^ vs. ln(number of particles). Thickness bins are defined as 100-125, 125-150, 150-175, 175-200, 200-225, 225-250, and >250 nm. Related to Fig. 1G.

**Supplementary Figure 4.**
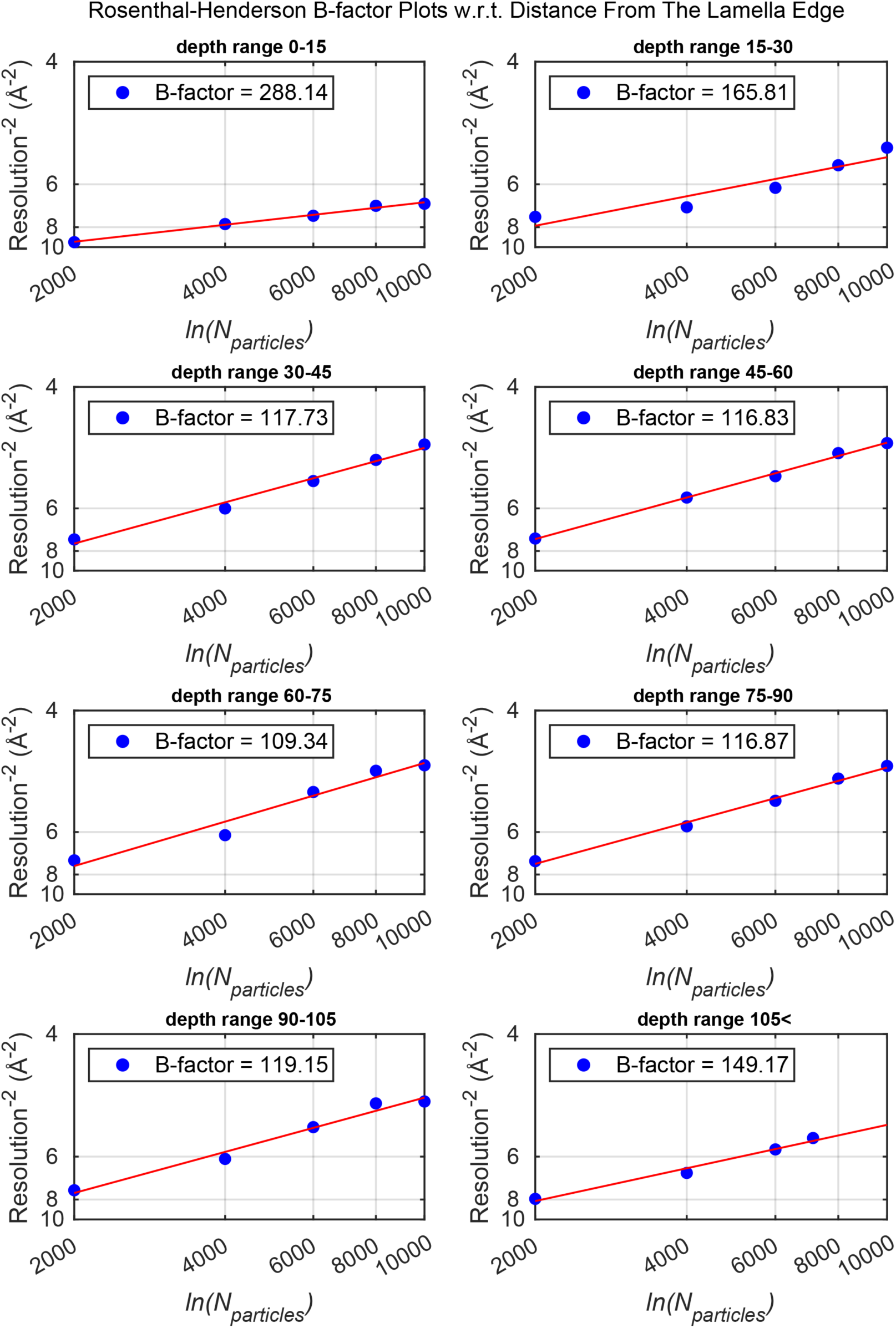
Rosenthal-Henderson B-factor plots with respect to depth from the lamella surface. B-factor is calculated based on the linear fit of resolution^−2^ vs. ln(number of particles). Depth bins are defined as 0-15, 15-30, 30-45, 45-60, 60-75, 75-90, 90-105, and >105 nm distance from the centers of ribosomes to the lamella surface. Related to Fig. 1H.

**Supplementary Figure 5.**
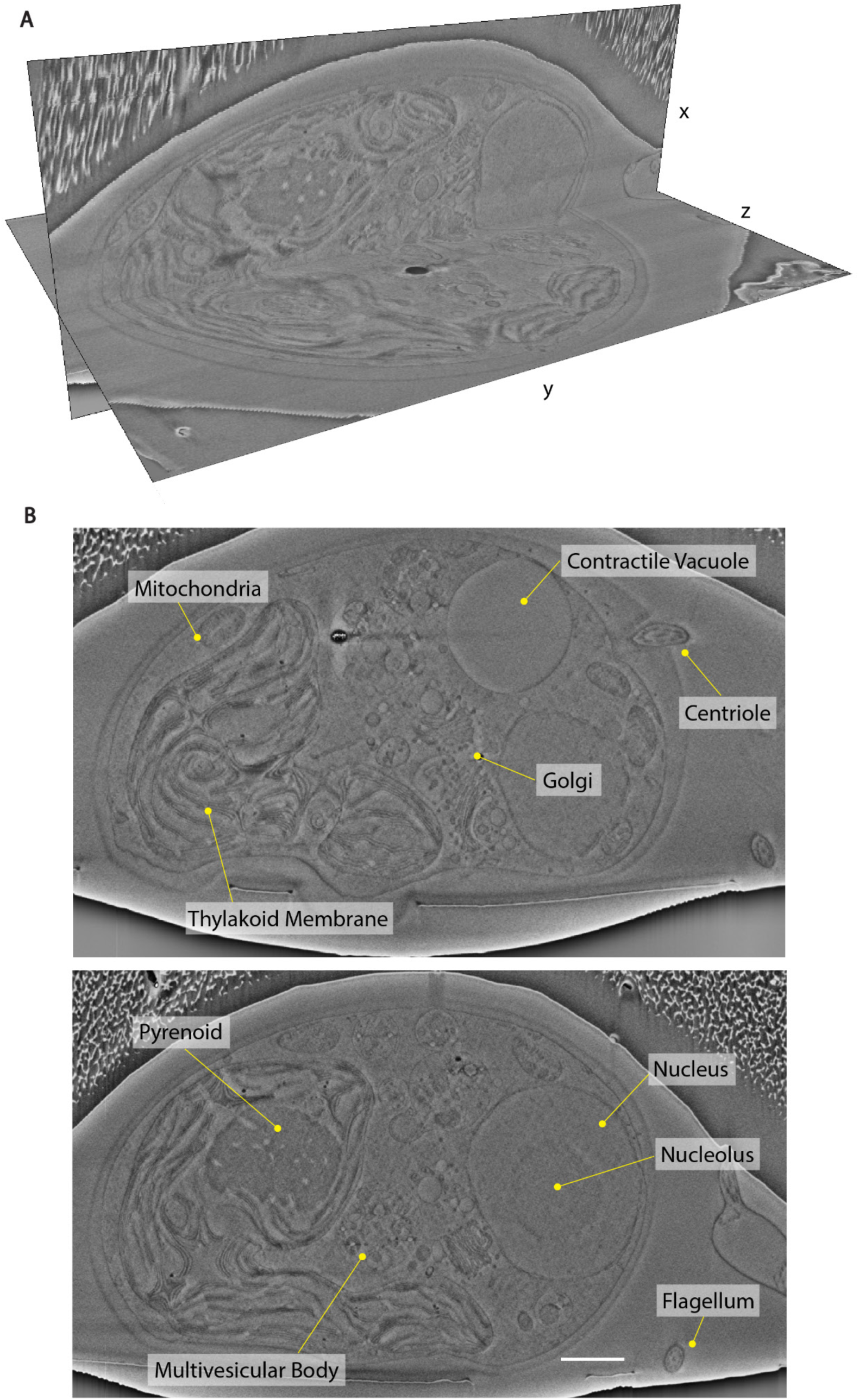
FIB-SEM slice and view data of a single *C. reinhardtii* cell (EMPIAR 11275). **A)** Two perpendicular cross section slices through the cell. Volumetric segmentation of this dataset was used for 3D visualization in Fig. 2A. **B)** Representative slices with visible organelles labelled. Scale bar: 1µm.

**Supplementary Figure 6.**
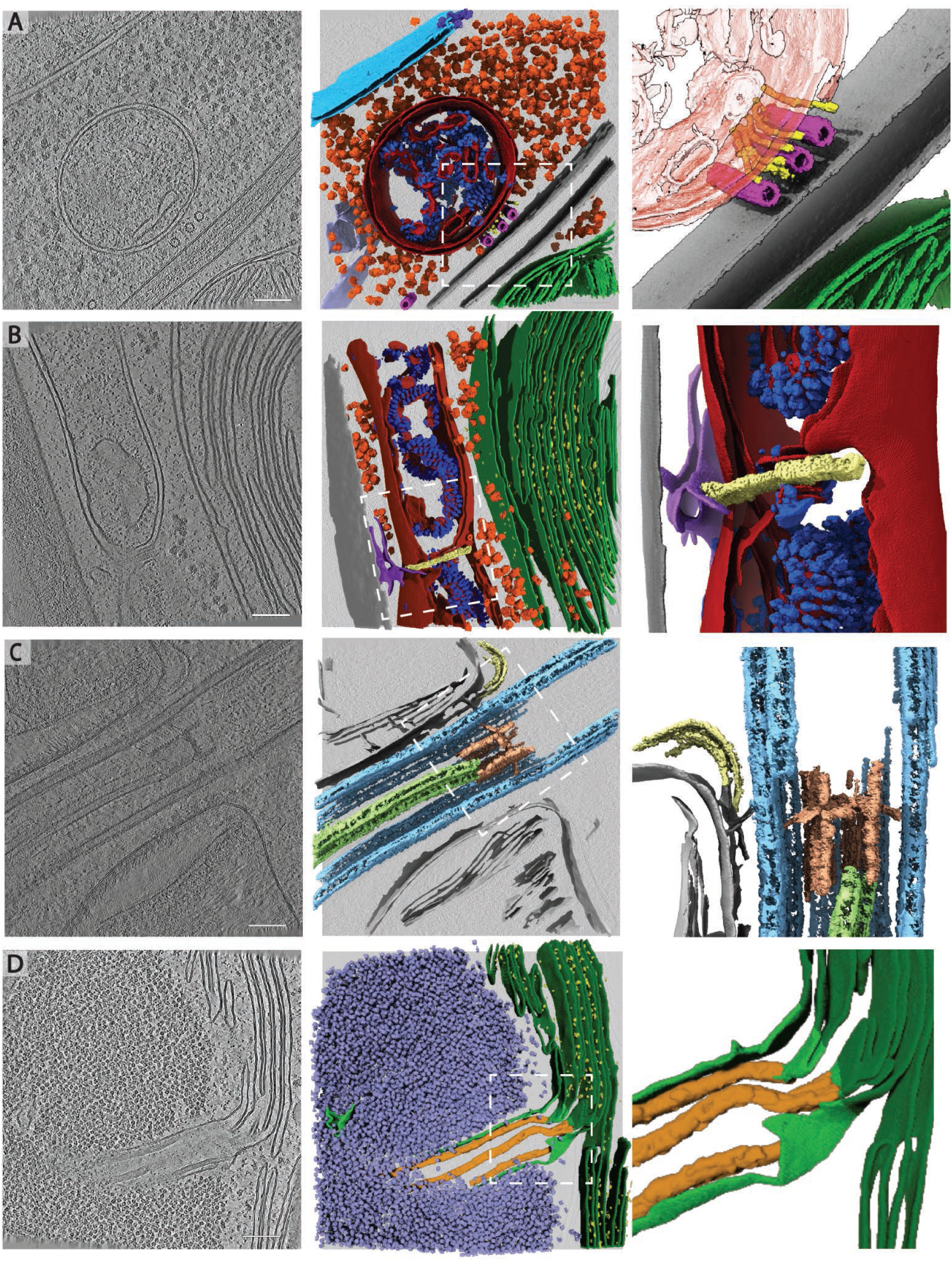
Rare cellular events observed in the cryo-ET dataset. Slices through tomograms (left) with corresponding 3D segmentation (middle and right). Right panels show enlarged views of the boxed regions in the middle panels. **A)** Mitochondrion at the plasma membrane, tethered to microtubules by thin filaments of unknown identity. Segmented classes: nuclear envelope (light blue), nuclear pore complex (dark blue), 80S ribosomes (orange), mitochondrial membranes (dark red), ATP synthase (royal blue), plasma membranes (grey), microtubules (pink), filaments (yellow), endoplasmic reticulum (purple). Zoomed-in panel shows thin filaments running between microtubules and the mitochondrion at the plasma membrane. Note that this tomogram contains two closely appressed cells, and thus, two plasma membranes with cell walls between. **B)** Mitochondrial fission event with ER membrane interactions. Segmented classes: mitochondrial membranes (dark red), ATP synthase (royal blue), 80S ribosomes (orange), thylakoid membranes (green), PSII (bright yellow), cell wall (grey), ER (purple), fission site containing filamentous structures perpendicular to the mitochondrial long axis (pale yellow). **C)** Ciliary transition zone between basal body and axoneme, including assembling IFT train and stellate structure^77^. Segmented classes: microtubule doublets (light blue), central microtubule pair (light green), stellate structure (pale red), IFT train (yellow). **D)** Pyrenoid tubule extending from the thylakoids into the phase-separated Rubisco matrix of the pyrenoid. Minitubules originate from thylakoid membranes^82^. Segmented classes: thylakoid membranes (dark green), PSII (yellow), pyrenoid tubule (lime green), Rubisco (lavender blue), minitubules (orange). Scale bars in A-D: 100 nm. Related to Fig. 2C-H.

**Supplementary Figure 7.**
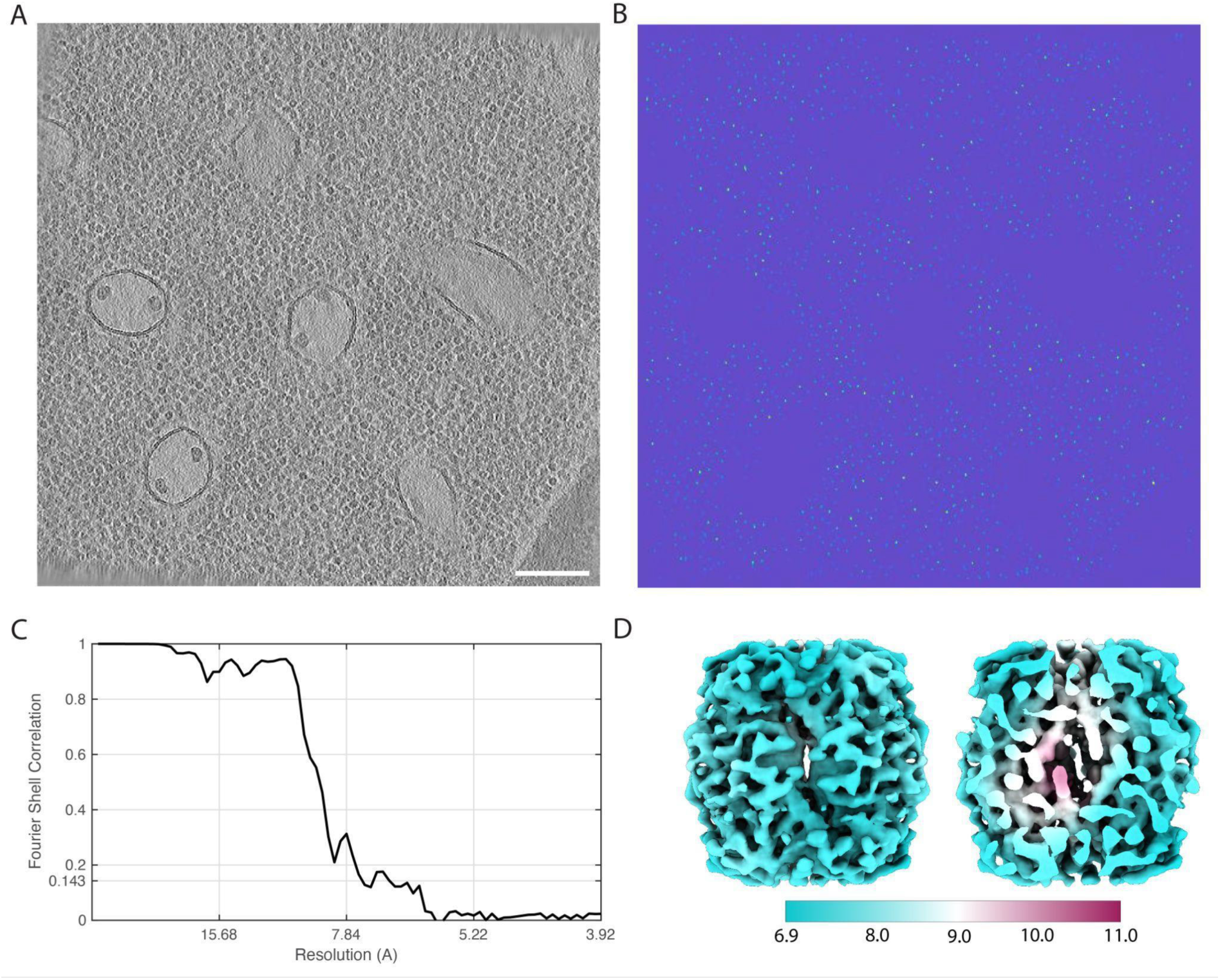
Supporting information for *in situ* STA of Rubisco. **A)** Slice through a tomogram of the pyrenoid, showing densely crowded Rubisco complexes. Scale bar: 100 nm. **B)** Corresponding slice through the output correlation volume after template matching with STOPGAP. Green and yellow pixels: higher cross-correlation scores, blue: low cross correlation. **C)** Global FSC plot (7.5 Å resolution) and **D)** local resolution map for the final STA map shown in Fig. 3C. Left: surface of the STA map, right: cross-section through the STA map.

**Supplementary Figure 8.**
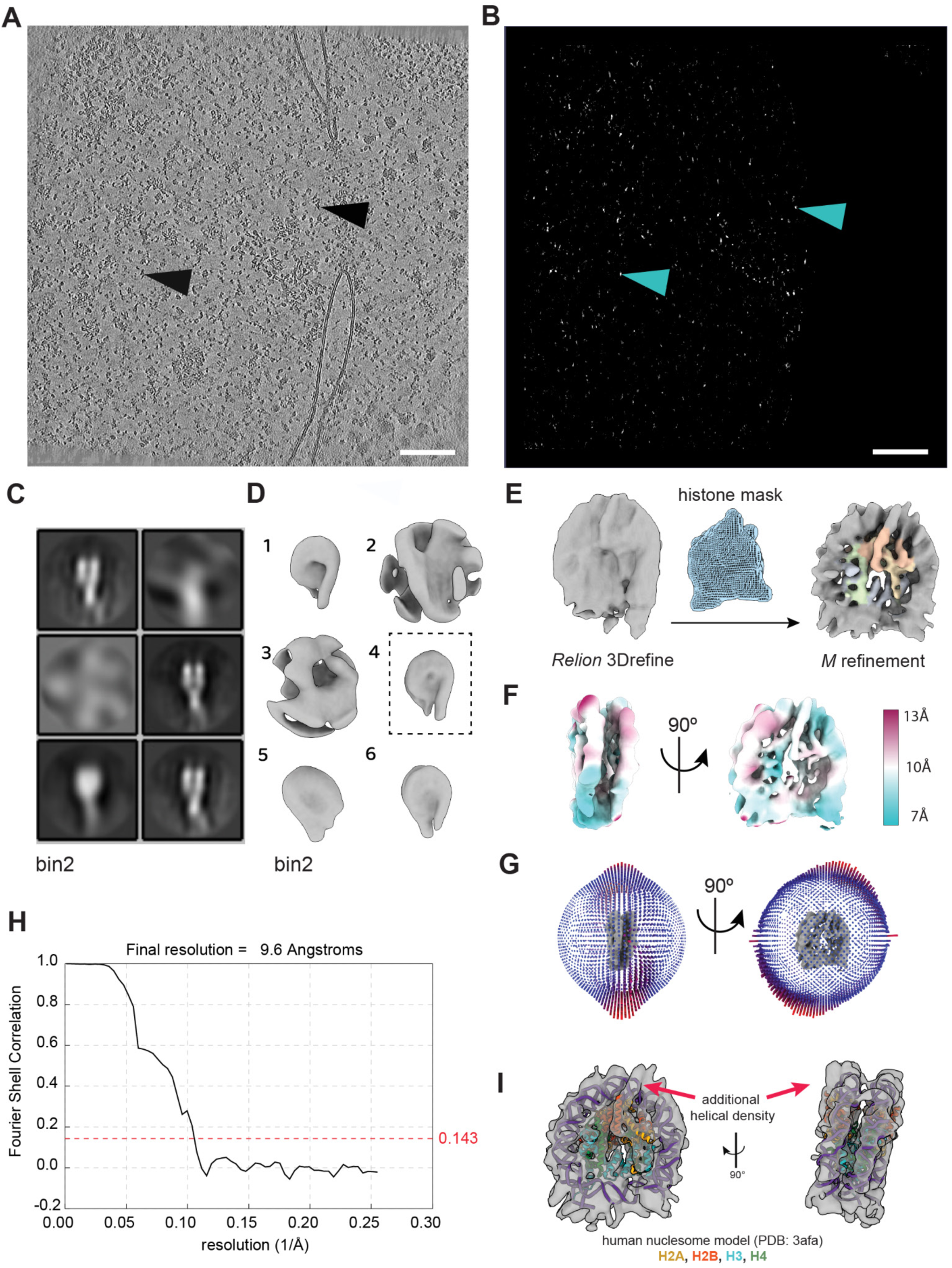
Supporting information for *in situ* STA of nucleosomes. **A)** Slice through a tomogram of the nucleus (left side), showing dense areas of chromatin, and cytoplasm outside of the nuclear envelope (right side). Black arrowheads indicate nucleosomes. **B)** Corresponding slice through the output correlation volume after template matching with PyTom-GPU^30^. Brighter pixels indicate higher cross-correlation results. Arrowheads indicate the correlation peak for the same nucleosome as in panel A. Scale bar in A-B: 100nm. **C-D)** Final 3D classification in RELION at 2x binning. **C)** Cross-section of each STA class and **D)** corresponding 3D maps. Particles within the class indicated in a dashed box were processed further in RELION and Warp/M. **E)** Following one round of 3D Refinement in RELION with a spherical mask, the resulting map and particles were imported into M and refined using a defined histone mask, resulting in the final average of 9.6 Å resolution obtained using ∼24k subtomograms. **F)** Local resolution, **G)** angular distribution, and **H)** FSC plot (9.6 Å resolution) for the final unbinned STA map shown in panel E and Fig. 3A. **I)** Fit of a human nucleosome atomic model (PDB: 3AFA)^141^into the STA map, revealing an extended H2B α-helix in *C. reinhardtii*.

**Supplementary Figure 9.**
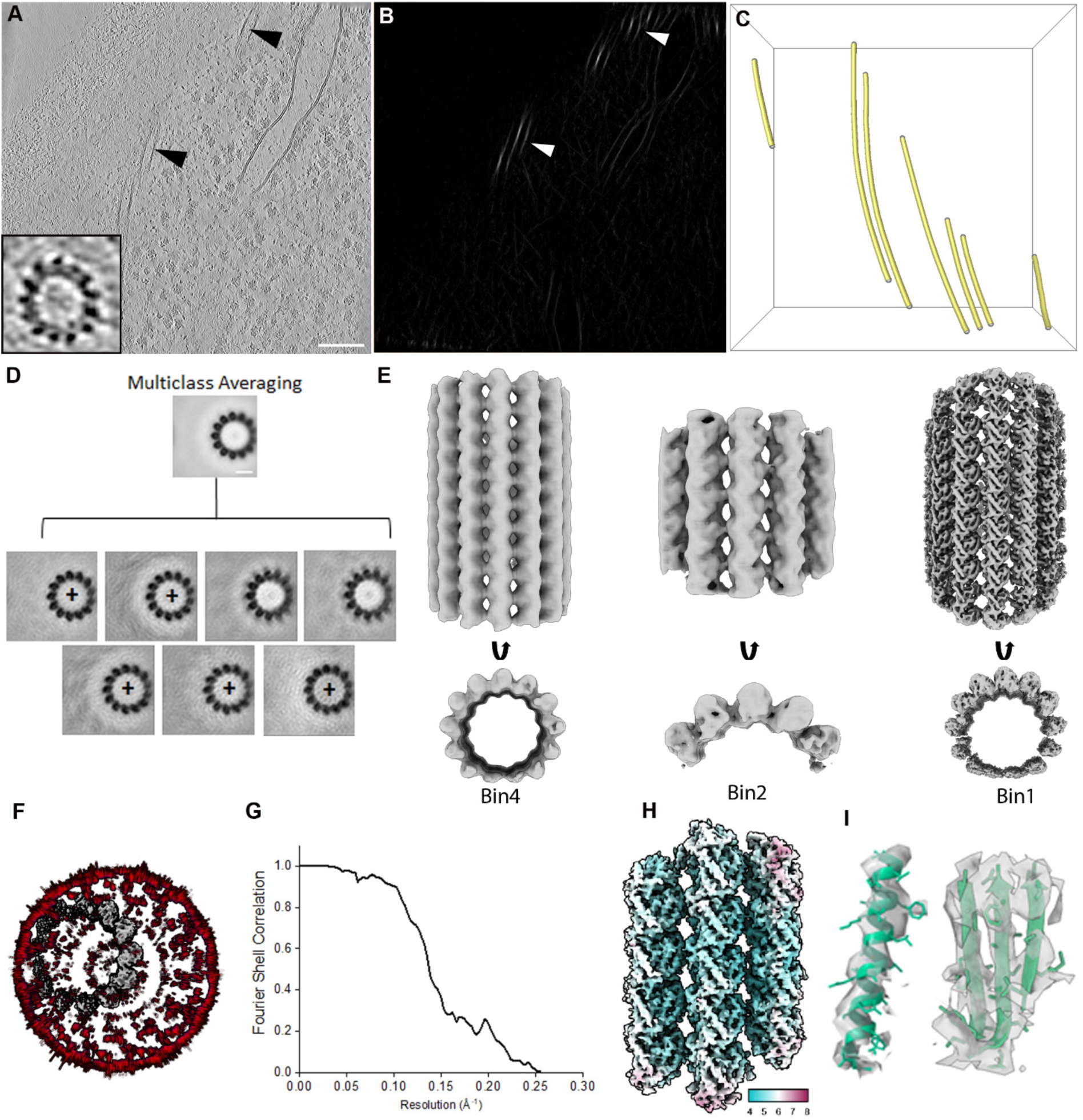
Supporting information for *in situ* STA of microtubules. **A)** Slice through a tomogram of the cytoplasm. Black arrowheads indicate MTs. Inset shows a zoom-in of a MT in cross-section, clearly resolving 13 protofilaments. Scale bar: 100 nm. **B**) Corresponding slice through the output correlation volume after template matching with Amira filament tracing. Brighter pixels indicate higher cross-correlation results. Arrowheads indicate the same MT as in panel A. **C)** 3D rendering of traced MT filaments (yellow). **D)** Multiclass averaging of the MTs for determination of the polarity from the radial skew of the tubulins as exemplified by plus sign. On a few occasions, MT polarity could not be determined unambiguously; therefore, these MTs were not included in the analysis. **E)** STA maps of MTs at 4x, 2x, and 1x binning (unbinned pixel size: 1.96 Å), obtained using ∼15k subtomograms. **F)** Angular distribution, **G**) FSC plot (4.7 Å resolution), and **H)** local resolution of the final unbinned STA map shown in Fig. 3D. **I)** Map quality depicted by rigid body fitting of a *C. reinhardtii* alpha-beta-tubulin dimer atomic model (green, PDB: 7JU4)^156^ into the STA map, showing representative fits of α-helix and β-sheet secondary structure elements as well as some bulky side chains.

**Supplementary Figure 10.**
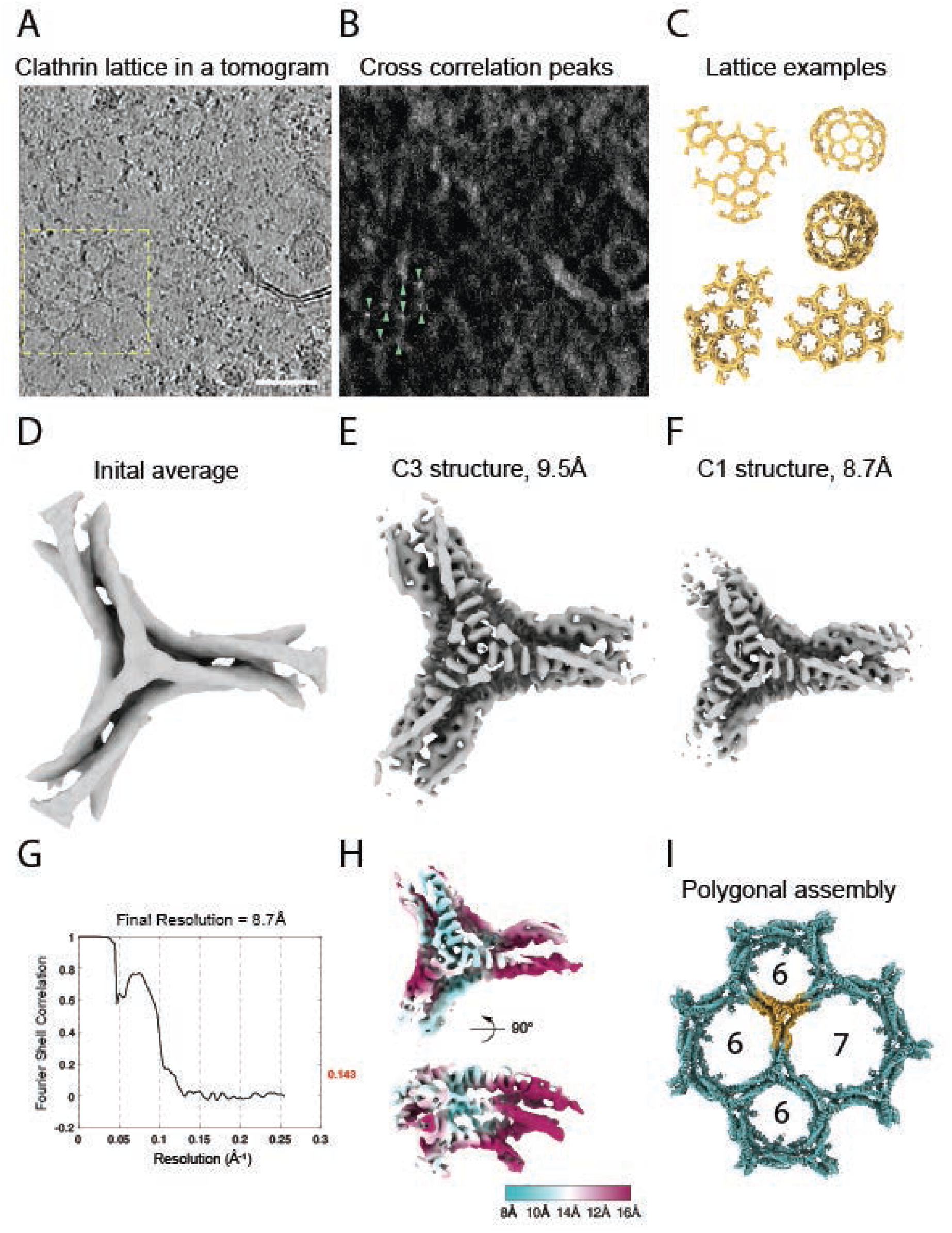
Supporting information for *in situ* STA of clathrin. **A)** Slice through a tomogram of the cytoplasm, showing an example of a clathrin lattice. Scale bar: 50 nm. **B)** Corresponding slice through the output correlation volume after template matching with PyTom. Brighter pixels indicate higher cross-correlation results. **C)** Examples of clathrin template placed back in a tomogram volume. Clathrin triskelia form characteristic lattice-like coats. **D)** Initial average from PyTom picks after cleaning the particle list. **E)** C3-symmetric clathrin structure at 9.5 Å resolution. **F)** Clathrin structure after symmetry expansion reaches 8.7 Å resolution. **G).** FSC plot (8.7 Å resolution) and **H)** local resolution of the clathrin STA map from panel F. **I)** Placing the STA map from panel E (orange) back into a tomogram reveals characteristic clathrin assemblies: pentagons, hexagons (also shown in Fig. 3E), and heptagons.

**Supplementary Figure 11.**
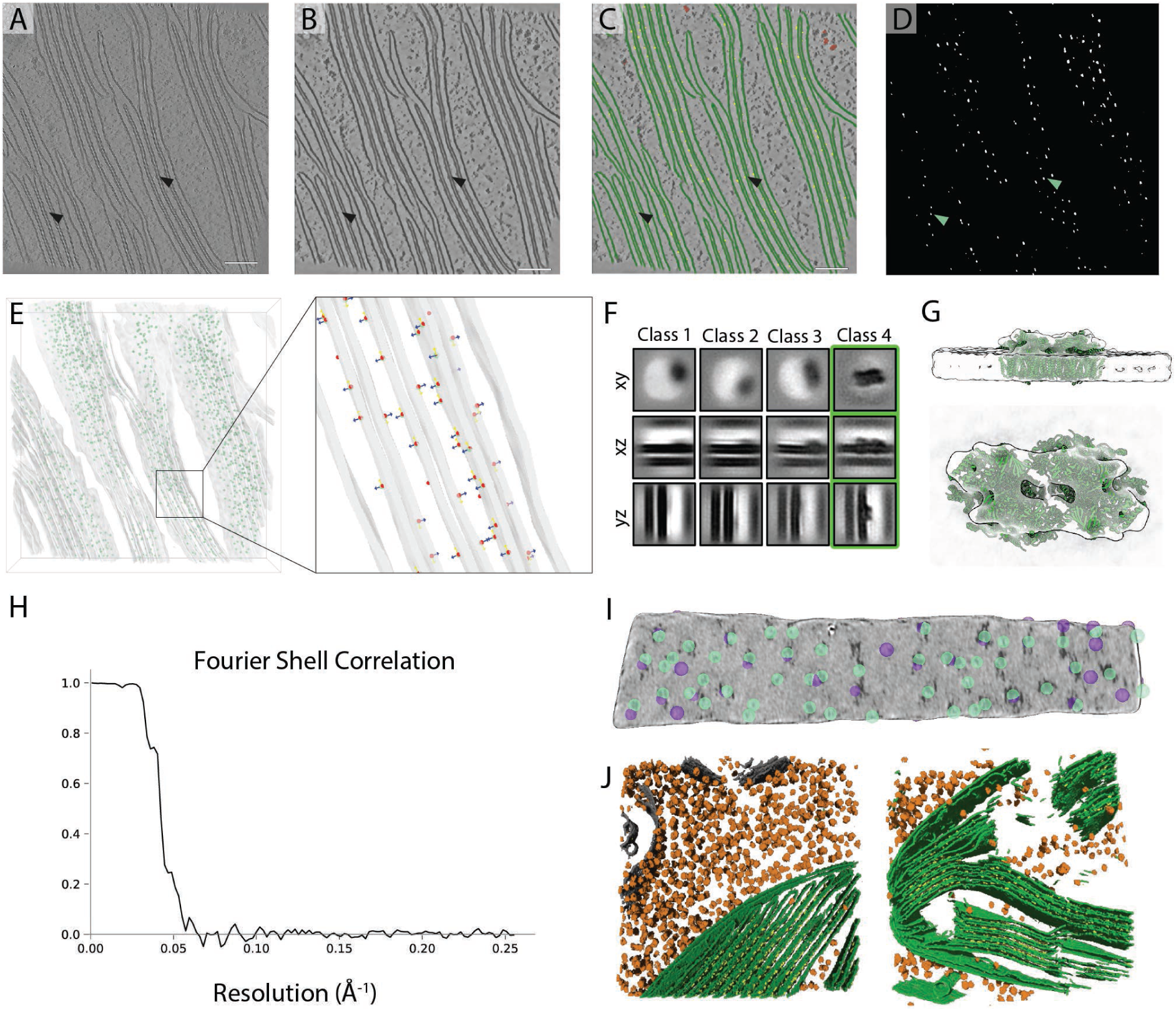
Supporting information for *in situ* STA of Photosystem II. In contrast to the other template-matching workflows, here a novel neural network picking scheme was developed. **A)** Representative tomogram of the chloroplast denoised with cryo-CARE^63^. Arrows point to PSII densities in thylakoid membranes. Scale bar: 100 nm. **B)** Regression U-net^88^ denoised tomogram. **C)** 2.5D semantic segmentation U-net trained with Dragonfly software to classify membrane (green), PSII (yellow), and ribosomes (orange). **D)** Binary mask of PSII candidate segmentations. **E)** Particle positions were extracted by centre of mass with initial angles acquired by normal assignment with regard to the membrane surface, using MemBrain^45^. Left: the PSII STA map placed back into the extracted positions. Right: zoom-in on positions with blue arrows indicating the normal vector. **F)** 2D projections of 3D class averages. **G)** Final STA map (from Fig. 3B) fit with a high-resolution structure (PDB: 6KAC)^109^, showing good agreement in the overall shape of PSII. **H)** Fourier Shell Correlation (FSC) plot of the unbinned subtomogram average obtained using 3,122 subtomograms (19 Å resolution). **I)** Comparison of template matching results (purple, not used for STA) and segmented coordinates (green, used for STA), overlaid on a “membranogram” showing PSII densities on the membrane surface. **J)** Representative 3D segmentations, with thylakoid membranes (green), PSII (yellow), and both 80S and chloroplast ribosomes (orange).

**Supplementary Figure 12.**
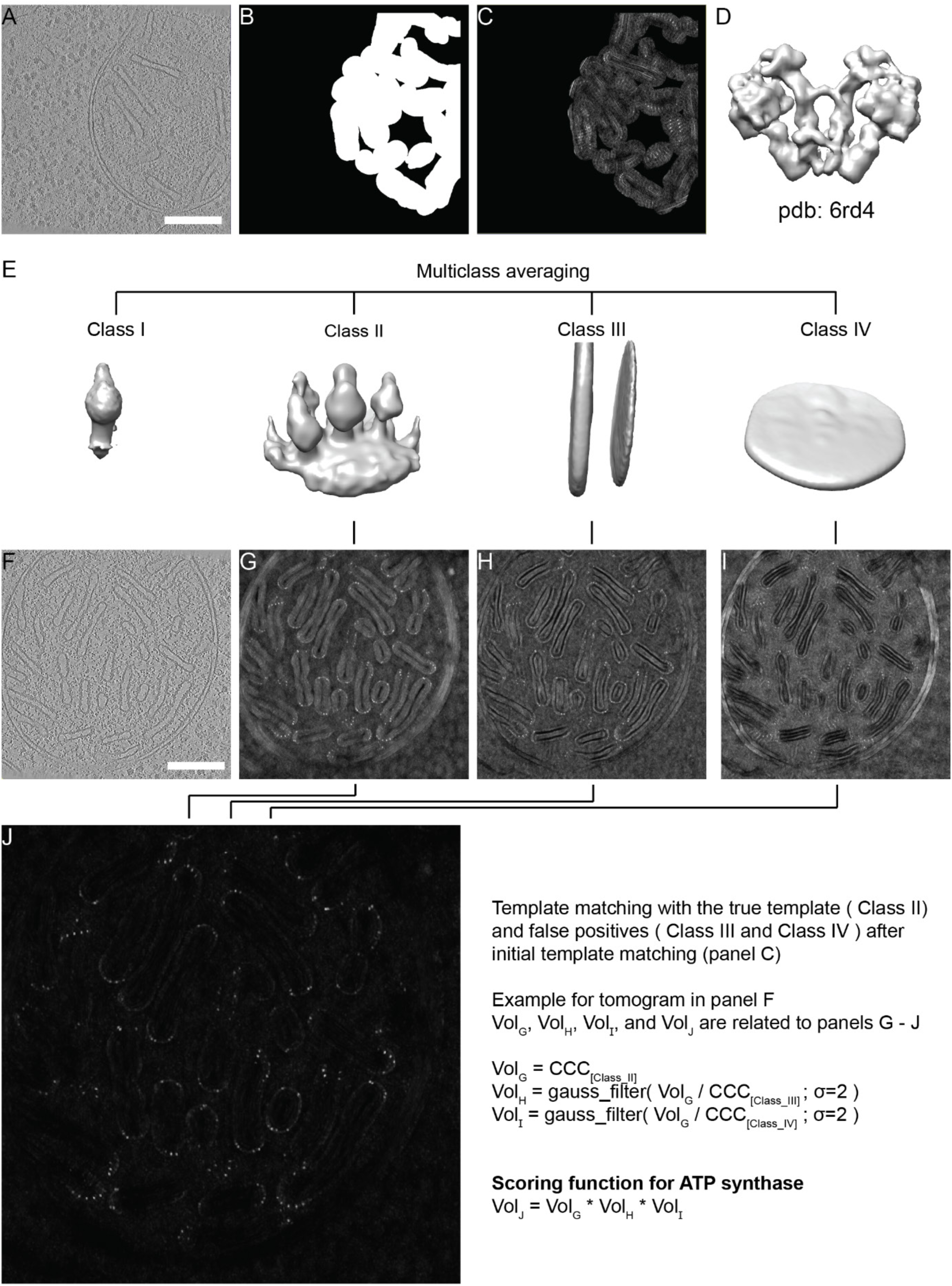
Supporting information for *in situ* STA of ATP synthase: template matching. **A)** Slice through a tomogram showing a mitochondrion in the cytoplasm. Scale bar: 200 nm. **B)** An expanded cristae mask used to select template matching peaks near crista membranes. **C)** Corresponding slice through the output correlation volume after initial template matching in STOPGAP and masking with panel B (masked score map). **D)** Density map simulated from PDB: 6RD4^113^ lowpass filtered to 15.7 Å, which was used for initial template matching. **E)** Class averages from multi-reference classification in STOPGAP (classes 1-4, from left to right). **F)** Slice through cryo-CARE-denoised tomogram – the score maps in panels G-J relate to the same tomogram. Scale bar: 200 nm. **G)** Template matching score map from PyTom-TM obtained for class 2 from panel E. **H,I)** Gaussian-filtered (σ=2) ratios of score map G and score maps obtained for classes 3 and 4, respectively. CCC: constrained cross-correlation. **J)** The final score map for picking subvolumes: the product of score maps G, H, and I. Brighter pixels indicate higher cross-correlation results. Related to Fig. 3F.

**Supplementary Figure 13.**
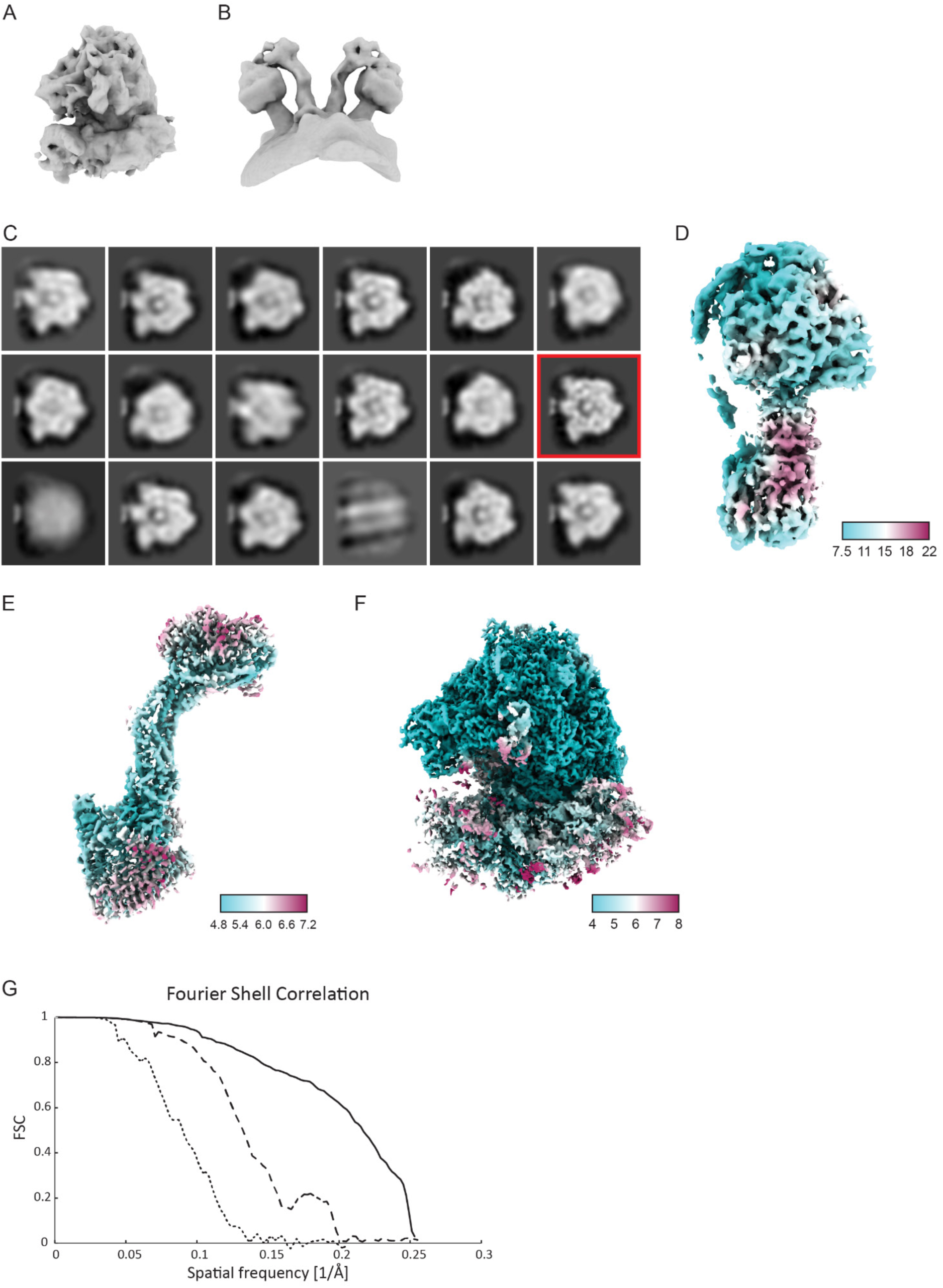
Supporting information for *in situ* STA of ATP synthase: subtomogram averaging and classification. **A, B)** 3D-refined 4x binned subtomogram averages for **A)** 80S ribosome and **B)** ATP synthase. **C)** 3D classification result showing different F_1_ head domain states. Each image shows a single slice through a different 3D class. **D-F)** Unbinned B-factor sharpened masked STA maps are shown with local resolution for: **D)** a selected class (marked by red outline in panel C) of ATP synthase capturing a single state of the head domain, as well as **E)** ATP synthase peripheral stalk and **F)** 80S consensus maps from multispecies refinement in M. **G)** FSC resolution estimates for the three unbinned STA maps: solid – 80S ribosome consensus map (4.0 Å), dashed – ATP synthase consensus map (5.2 Å), dotted – map of ATP synthase classified state (8.6 Å). STA maps from panels D and E were combined to make the composite map shown in Fig. 3F.

**Supplementary Figure 14.**
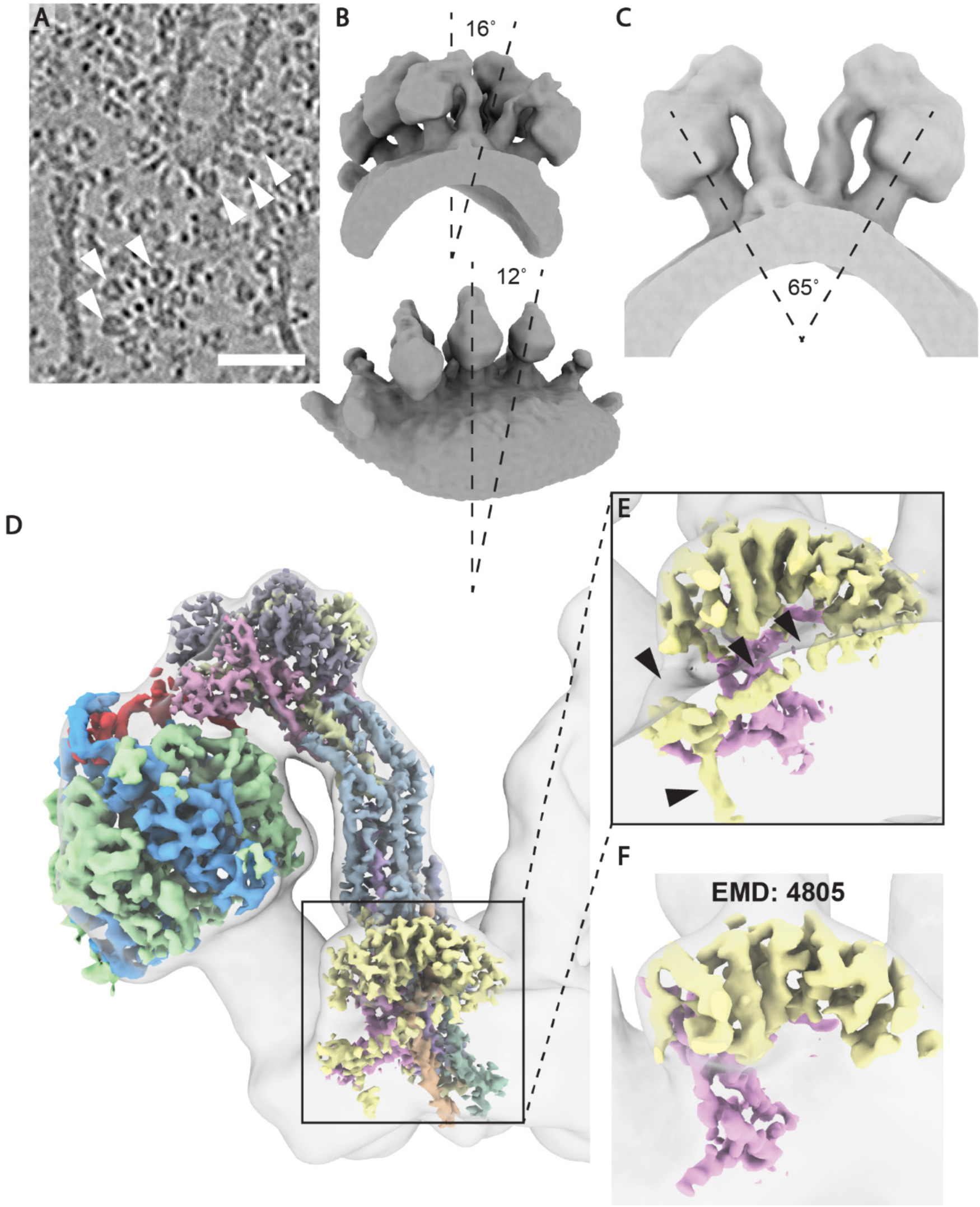
Supporting information for *in situ* STA of ATP synthase: details of STA map. **A)** Tomogram zoom-in showing cristae decorated with ATP synthase dimers. Examples are indicated by white arrowheads on a tomogram slice. Scale bar: 50 nm. **B)** Front and side view of ATP synthase 4x binned STA map showing three synthase dimers forming a row. Twist (16°) and (12°) bend angles are indicated by dashed lines. **C)** Cut-through ATP synthase STA map, with 65° angle between the two central stalks marked by dashed lines. **E)** Unbinned composite map consisting of a refined consensus map showing individual subunits from the peripheral stalk and a refined map of a single state of the head domain (α-subunit – blue; β-subunit – light green) obtained by 3D classification. This 1x binned composite map (colors) is shown fit within the 4x binned map from panel C (grey). **E)** Inset from panel D, highlighting the a-subunit (pink) and ASA3 (yellow). Densities that have not been identified previously by SPA are indicated by black arrowheads. **F)** ASA3 (yellow) and the a-subunit (pink) of *Polytomella sp.* segmented out of the EMDB-4805 SPA map low-pass filtered to 5 Å. Note the differences compared to panel E.

**Supplementary Figure 15.**
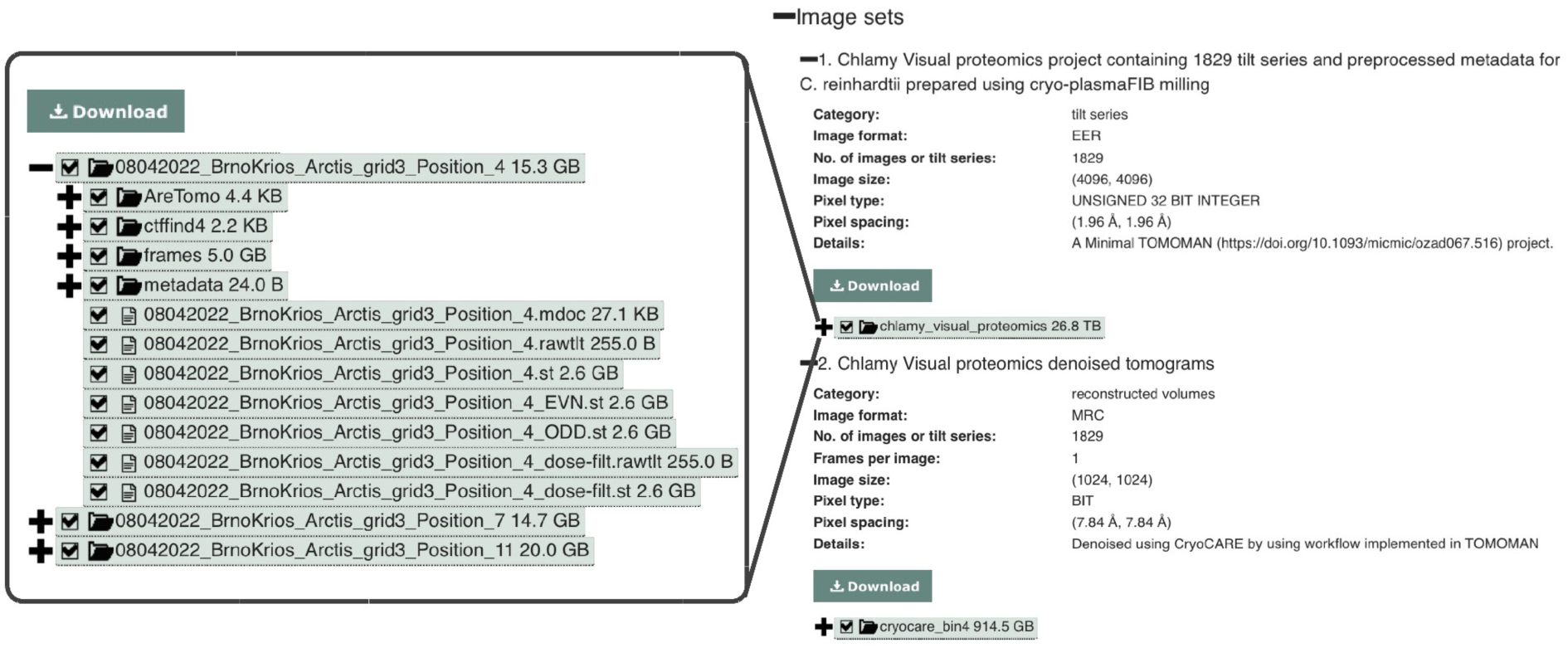
Data structure of EMPIAR-11830. Two image sets were created, with the first one containing all tilt-series. Each tilt-series is split into a separate subfolder accommodating a separate minimal TOMOMAN project. The second image set presents all reconstructed tomograms numbered chronologically respectively to their tilt-series.

## References

1. Nickell, S., Kofler, C., Leis, A. P. & Baumeister, W. A visual approach to proteomics. Nat. Rev. Mol. Cell Biol. 7, 225–230 (2006).

2. McCafferty, C. L. et al. Integrating cellular electron microscopy with multimodal data to explore biology across space and time. Cell 187, 563–584 (2024).

3. Young, L. N. & Villa, E. Bringing Structure to Cell Biology with Cryo-Electron Tomography. Annu. Rev. Biophys. 52, 573–595 (2023).

4. Nogales, E. & Mahamid, J. Bridging structural and cell biology with cryo-electron microscopy. Nature 628, 47–56 (2024).

5. Ferreira, J. L. et al. Variable microtubule architecture in the malaria parasite. Nat. Commun. 14, 1216 (2023).

6. Rodrigues-Oliveira, T. et al. Actin cytoskeleton and complex cell architecture in an Asgard archaeon. Nature 613, 332–339 (2023).

7. Dietrich, H. M. et al. Membrane-anchored HDCR nanowires drive hydrogen-powered CO2 fixation. Nature 607, 823–830 (2022).

8. Jasnin, M. et al. The Architecture of Traveling Actin Waves Revealed by Cryo-Electron Tomography. Structure 27, 1211–1223.e5 (2019).

9. Zimmerli, C. E. et al. Nuclear pores dilate and constrict in cellulo. Science 374, eabd9776 (2021).

10. Tamborrini, D. et al. Structure of the native myosin filament in the relaxed cardiac sarcomere. Nature 623, 863–871 (2023).

11. Pöge, M. et al. Determinants shaping the nanoscale architecture of the mouse rod outer segment. Elife 10, (2021).

12. Laughlin, T. G. et al. Architecture and self-assembly of the jumbo bacteriophage nuclear shell. Nature 608, 429–435 (2022).

13. Schiøtz, O. H. et al. Serial Lift-Out: sampling the molecular anatomy of whole organisms. Nat. Methods (2023) doi:10.1038/s41592-023-02113-5.

14. Zens, B. et al. Lift-out cryo-FIBSEM and cryo-ET reveal the ultrastructural landscape of extracellular matrix. J. Cell Biol. 223, (2024).

15. Creekmore, B. C., Kixmoeller, K., Black, B. E., Lee, E. B. & Chang, Y.-W. Ultrastructure of human brain tissue vitrified from autopsy revealed by cryo-ET with cryo-plasma FIB milling. Nat. Commun. 15, 2660 (2024).

16. Klumpe, S. et al. A modular platform for automated cryo-FIB workflows. Elife 10, (2021).

17. Cleeve, P. et al. OpenFIBSEM: A universal API for FIBSEM control. J. Struct. Biol. 215, 107967 (2023).

18. Zachs, T. et al. Fully automated, sequential focused ion beam milling for cryo-electron tomography. Elife 9, (2020).

19. Tacke, S. et al. A streamlined workflow for automated cryo focused ion beam milling. J. Struct. Biol. 213, 107743 (2021).

20. Khavnekar, S. et al. Multishot tomography for high-resolution in situ subtomogram averaging. J. Struct. Biol. 215, 107911 (2023).

21. Bouvette, J. et al. Beam image-shift accelerated data acquisition for near-atomic resolution single-particle cryo-electron tomography. Nat. Commun. 12, 1957 (2021).

22. Eisenstein, F. et al. Parallel cryo electron tomography on in situ lamellae. Nat. Methods 20, 131–138 (2023).

23. Eisenstein, F., Fukuda, Y. & Danev, R. Smart parallel automated cryo-electron tomography. Nat. Methods 21, 1612–1615 (2024).

24. Yang, J. E. et al. Correlative montage parallel array cryo-tomography for in situ structural cell biology. Nat. Methods 20, 1537–1543 (2023).

25. Berger, C. et al. Plasma FIB milling for the determination of structures in situ. Nat. Commun. 14, 629 (2023).

26. Smith, N. et al. High brightness inductively coupled plasma source for high current focused ion beam applications. J. Vac. Sci. Technol. B Microelectron. Nanometer Struct. Process. Meas. Phenom. 24, 2902–2906 (2006).

27. Cruz-León, S. et al. High-confidence 3D template matching for cryo-electron tomography. Nat. Commun. 15, 3992 (2024).

28. Lucas, B. A., Himes, B. A. & Grigorieff, N. Baited reconstruction with 2D template matching for high-resolution structure determination in vitro and in vivo without template bias. Elife 12, (2023).

29. Wan, W., Khavnekar, S. & Wagner, J. STOPGAP: an open-source package for template matching, subtomogram alignment and classification. Acta Crystallogr D Struct Biol (2024) doi:10.1107/S205979832400295X.

30. Hrabe, T. et al. PyTom: a python-based toolbox for localization of macromolecules in cryo-electron tomograms and subtomogram analysis. J. Struct. Biol. 178, 177–188 (2012).

31. Chaillet, M. L. et al. Extensive Angular Sampling Enables the Sensitive Localization of Macromolecules in Electron Tomograms. Int. J. Mol. Sci. 24, (2023).

32. Tegunov, D., Xue, L., Dienemann, C., Cramer, P. & Mahamid, J. Multi-particle cryo-EM refinement with M visualizes ribosome-antibiotic complex at 3.5 Å in cells. Nat. Methods 18, 186–193 (2021).

33. Burt, A. et al. An image processing pipeline for electron cryo-tomography in RELION-5. bioRxiv 2024.04.26.591129 (2024) doi:10.1101/2024.04.26.591129.

34. Fäßler, F., Dimchev, G., Hodirnau, V.-V., Wan, W. & Schur, F. K. M. Cryo-electron tomography structure of Arp2/3 complex in cells reveals new insights into the branch junction. Nat. Commun. 11, 6437 (2020).

35. Zhang, X. et al. Molecular mechanisms of stress-induced reactivation in mumps virus condensates. Cell 186, 1877–1894.e27 (2023).

36. Sutton, G. et al. Assembly intermediates of orthoreovirus captured in the cell. Nat. Commun. 11, 4445 (2020).

37. Wang, Z. et al. Structures from intact myofibrils reveal mechanism of thin filament regulation through nebulin. Science 375, eabn1934 (2022).

38. Xue, L. et al. Visualizing translation dynamics at atomic detail inside a bacterial cell. Nature 610, 205–211 (2022).

39. Xing, H. et al. Translation dynamics in human cells visualized at high resolution reveal cancer drug action. Science 381, 70–75 (2023).

40. Hoffmann, P. C. et al. Structures of the eukaryotic ribosome and its translational states in situ. Nat. Commun. 13, 7435 (2022).

41. Anton, L. et al. Multiscale effects of perturbed translation dynamics inform antimalarial design. bioRxiv 2023.09.03.556115 (2023) doi:10.1101/2023.09.03.556115.

42. Zheng, W., et al. Visualizing the translation landscape in human cells at high resolution. bioRxiv (2024) doi:10.1101/2024.07.02.601723.

43. Rickgauer, J. P., Choi, H., Moore, A. S., Denk, W. & Lippincott-Schwartz, J. Structural dynamics of human ribosomes in situ reconstructed by exhaustive high-resolution template matching. Mol. Cell (2024) doi:10.1016/j.molcel.2024.11.003.

44. Rice, G. et al. TomoTwin: generalized 3D localization of macromolecules in cryo-electron tomograms with structural data mining. Nat. Methods 20, 871–880 (2023).

45. Lamm, L., et al. MemBrain v2: an end-to-end tool for the analysis of membranes in cryo-electron tomography. bioRxiv 2024.01.05.574336 (2024) doi:10.1101/2024.01.05.574336.

46. de Teresa-Trueba, I. et al. Convolutional networks for supervised mining of molecular patterns within cellular context. Nat. Methods 20, 284–294 (2023).

47. Moebel, E. et al. Deep learning improves macromolecule identification in 3D cellular cryo-electron tomograms. Nat. Methods 18, 1386–1394 (2021).

48. Chen, M. et al. Convolutional neural networks for automated annotation of cellular cryo-electron tomograms. Nat. Methods 14, 983–985 (2017).

49. Iudin, A. et al. EMPIAR: the Electron Microscopy Public Image Archive. Nucleic Acids Res. 51, D1503–D1511 (2023).

50. Ermel, U. et al. A data portal for providing standardized annotations for cryo-electron tomography. Nat. Methods 21, 2200–2202 (2024).

51. Goodenough, U. & Engel, B. D. Chapter 2 - Cell ultrastructure. in The Chlamydomonas Sourcebook (Third *Edition)* (ed. Goodenough, U.) 17–40 (Academic Press, 2023). doi:10.1016/B978-0-12-822457-1.00015-7.

52. Jinkerson, R. E. & Jonikas, M. C. Molecular techniques to interrogate and edit the Chlamydomonas nuclear genome. Plant J. 82, 393–412 (2015).

53. Crozet, P. et al. Birth of a Photosynthetic Chassis: A MoClo Toolkit Enabling Synthetic Biology in the Microalga Chlamydomonas reinhardtii. ACS Synth. Biol. 7, 2074–2086 (2018).

54. Salomé, P. A. & Merchant, S. S. A Series of Fortunate Events: Introducing Chlamydomonas as a Reference Organism. Plant Cell 31, 1682–1707 (2019).

55. Sasso, S., Stibor, H., Mittag, M. & Grossman, A. R. From molecular manipulation of domesticated Chlamydomonas reinhardtii to survival in nature. Elife 7, (2018).

56. Marshall, W. F. Chlamydomonas as a model system to study cilia and flagella using genetics, biochemistry, and microscopy. Front. Cell Dev. Biol. 12, (2024).

57. Sergey, G., et al. Oxygen plasma focused ion beam scanning electron microscopy for biological samples. bioRxiv 457820 (2018) doi:10.1101/457820.

58. Kuba, J. et al. Advanced cryo-tomography workflow developments - correlative microscopy, milling automation and cryo-lift-out. J. Microsc. 281, 112–124 (2021).

59. Brogden, V. et al. Material Sputtering with a Multi-Ion Species Plasma Focused Ion Beam. Advances in Materials Science and Engineering 2021, (2021).

60. Dumoux, M. et al. Cryo-plasma FIB/SEM volume imaging of biological specimens. Elife 12, (2023).

61. Berger, C., Watson, H., Naismith, J., Dumoux, M. & Grange, M. Xenon plasma focused ion beam lamella fabrication on high-pressure frozen specimens for structural cell biology. bioRxiv (2024) doi:10.1101/2024.06.20.599830.

62. Spurný, R. et al. Arctis WebUI - A Novel Software Concept for Automating Cryo-lamellae Production. Microsc. Microanal. 29, 2081–2082 (2023).

63. Buchholz, T.-O., Jordan, M., Pigino, G. & Jug, F. Cryo-CARE: Content-Aware Image Restoration for Cryo-Transmission Electron Microscopy Data. in 2019 IEEE 16th International Symposium on Biomedical Imaging (ISBI 2019) 502–506 (IEEE, 2019). doi:10.1109/ISBI.2019.8759519.

64. Rangan, R. et al. CryoDRGN-ET: deep reconstructing generative networks for visualizing dynamic biomolecules inside cells. Nat. Methods (2024) doi:10.1038/s41592-024-02340-4.

65. Pfeffer, S. et al. Dissecting the molecular organization of the translocon-associated protein complex. Nat. Commun. 8, 14516 (2017).

66. Tuijtel, M. W. et al. Thinner is not always better: Optimizing cryo-lamellae for subtomogram averaging. Sci Adv 10, eadk6285 (2024).

67. Lucas, B. A. & Grigorieff, N. Quantification of gallium cryo-FIB milling damage in biological lamellae. Proc. Natl. Acad. Sci. U. S. A. 120, e2301852120 (2023).

68. Albert, S. et al. Proteasomes tether to two distinct sites at the nuclear pore complex. Proc. Natl. Acad. Sci. U. S. A. 114, 13726–13731 (2017).

69. Erdmann, P. S. et al. In situ cryo-electron tomography reveals gradient organization of ribosome biogenesis in intact nucleoli. Nat. Commun. 12, 5364 (2021).

70. Mosalaganti, S. et al. In situ architecture of the algal nuclear pore complex. Nat. Commun. 9, 2361 (2018).

71. Albert, S. et al. Direct visualization of degradation microcompartments at the ER membrane. Proc. Natl. Acad. Sci. U. S. A. 117, 1069–1080 (2020).

72. Engel, B. D. et al. In situ structural analysis of Golgi intracisternal protein arrays. Proc. Natl. Acad. Sci. U. S. A. 112, 11264–11269 (2015).

73. Bykov, Y. S. et al. The structure of the COPI coat determined within the cell. Elife 6, (2017).

74. Kovtun, O. et al. Structure of the membrane-assembled retromer coat determined by cryo-electron tomography. Nature 561, 561–564 (2018).

75. Le Guennec, M. et al. A helical inner scaffold provides a structural basis for centriole cohesion. Sci Adv 6, eaaz4137 (2020).

76. Klena, N. et al. Architecture of the centriole cartwheel-containing region revealed by cryo-electron tomography. EMBO J. 39, e106246 (2020).

77. van den Hoek, H., et al. In situ architecture of the ciliary base reveals the stepwise assembly of intraflagellar transport trains. Science 377, 543–548 (2022).

78. Jordan, M. A., Diener, D. R., Stepanek, L. & Pigino, G. The cryo-EM structure of intraflagellar transport trains reveals how dynein is inactivated to ensure unidirectional anterograde movement in cilia. Nat. Cell Biol. 20, 1250–1255 (2018).

79. Lacey, S. E., Foster, H. E. & Pigino, G. The molecular structure of IFT-A and IFT-B in anterograde intraflagellar transport trains. Nat. Struct. Mol. Biol. 30, 584–593 (2023).

80. Craig, E. W. et al. The elusive actin cytoskeleton of a green alga expressing both conventional and divergent actins. Mol. Biol. Cell 30, 2827–2837 (2019).

81. Waltz, F. et al. How to build a ribosome from RNA fragments in Chlamydomonas mitochondria. Nat. Commun. 12, 7176 (2021).

82. Engel, B. D. et al. Native architecture of the Chlamydomonas chloroplast revealed by in situ cryo-electron tomography. Elife 4, (2015).

83. Freeman Rosenzweig, E. S., et al. The Eukaryotic CO2-Concentrating Organelle Is Liquid-like and Exhibits Dynamic Reorganization. Cell 171, 148–162.e19 (2017).

84. Wietrzynski, W. et al. Charting the native architecture of Chlamydomonas thylakoid membranes with single-molecule precision. Elife 9, (2020).

85. Wietrzynski, W. & Engel, B. D. Chapter 23 - Supramolecular organization of chloroplast membranes. in The Chlamydomonas Sourcebook (Third *Edition)* (eds. Grossman, A. R. & Wollman, F.-A.) 763–785 (Academic Press, London, 2023). doi:10.1016/B978-0-12-821430-5.00018-3.

86. Peddie, C. J. et al. Volume electron microscopy. Nat Rev Methods Primers 2, 51 (2022).

87. Xu, C. S. et al. Enhanced FIB-SEM systems for large-volume 3D imaging. Elife 6, (2017).

88. Purnell, C., et al. Rapid Synthesis of Cryo-ET Data for Training Deep Learning Models. bioRxiv (2023) doi:10.1101/2023.04.28.538636.

89. Heebner, J. E. et al. Deep Learning-Based Segmentation of Cryo-Electron Tomograms. J. Vis. Exp. (2022) doi:10.3791/64435.

90. Friedman, J. R. et al. ER tubules mark sites of mitochondrial division. Science 334, 358–362 (2011).

91. Ohad, I., Siekevitz, P. & Palade, G. E. Biogenesis of chloroplast membranes. I. Plastid dedifferentiation in a dark-grown algal mutant (Chlamydomonas reinhardi). J. Cell Biol. 35, 521–552 (1967).

92. Hennacy, J. H. et al. SAGA1 and MITH1 produce matrix-traversing membranes in the CO2-fixing pyrenoid. Nat. Plants (2024) doi:10.1038/s41477-024-01847-0.

93. Franklin, E., et al. Proteomic analysis of the pyrenoid-traversing membranes of*Chlamydomonas reinhardtii* reveals novel components. bioRxiv (2024) doi:10.1101/2024.10.28.620638.

94. Taylor, T. C., Backlund, A., Bjorhall, K., Spreitzer, R. J. & Andersson, I. First crystal structure of Rubisco from a green alga, Chlamydomonas reinhardtii. J. Biol. Chem. 276, 48159–48164 (2001).

95. Luger, K., Mäder, A. W., Richmond, R. K., Sargent, D. F. & Richmond, T. J. Crystal structure of the nucleosome core particle at 2.8 A resolution. Nature 389, 251–260 (1997).

96. Fatmaoui, F. et al. Cryo-electron tomography and deep learning denoising reveal native chromatin landscapes of interphase nuclei. bioRxiv 2022.08.16.502515 (2022) doi:10.1101/2022.08.16.502515.

97. Cai, S., Böck, D., Pilhofer, M. & Gan, L. The in situ structures of mono-, di-, and trinucleosomes in human heterochromatin. Mol. Biol. Cell 29, 2450–2457 (2018).

98. Chen, J. K. et al. Nanoscale analysis of human G1 and metaphase chromatin in situ. bioRxiv 2023.07.31.551204 (2023) doi:10.1101/2023.07.31.551204.

99. Tan, Z. Y. et al. Heterogeneous non-canonical nucleosomes predominate in yeast cells in situ. Elife 12, (2023).

100. Hou, Z., Nightingale, F., Zhu, Y., MacGregor-Chatwin, C. & Zhang, P. Structure of native chromatin fibres revealed by Cryo-ET in situ. Nat. Commun. 14, 6324 (2023).

101. Mittelmeier, T. M., Boyd, J. S., Lamb, M. R. & Dieckmann, C. L. Asymmetric properties of the Chlamydomonas reinhardtii cytoskeleton direct rhodopsin photoreceptor localization. J. Cell Biol. 193, 741–753 (2011).

102. Boyd, J. S., Gray, M. M., Thompson, M. D., Horst, C. J. & Dieckmann, C. L. The daughter four-membered microtubule rootlet determines anterior-posterior positioning of the eyespot in Chlamydomonas reinhardtii. Cytoskeleton 68, 459–469 (2011).

103. Tilney, L. G. et al. Microtubules: evidence for 13 protofilaments. J. Cell Biol. 59, 267–275 (1973).

104. Chalfie, M. & Thomson, J. N. Structural and functional diversity in the neuronal microtubules of Caenorhabditis elegans. J. Cell Biol. 93, 15–23 (1982).

105. Chakraborty, S., et al. Cryo-electron tomography suggests tubulin chaperones form a subset of microtubule lumenal particles with a role in maintaining neuronal microtubules. bioRxiv 2022.07.28.501854 (2024) doi:10.1101/2022.07.28.501854.

106. Foster, H. E., Ventura Santos, C. & Carter, A. P. A cryo-ET survey of microtubules and intracellular compartments in mammalian axons. J. Cell Biol. 221, (2022).

107. Morris, K. L. et al. Cryo-EM of multiple cage architectures reveals a universal mode of clathrin self-assembly. Nat. Struct. Mol. Biol. 26, 890–898 (2019).

108. Serwas, D. et al. Mechanistic insights into actin force generation during vesicle formation from cryo-electron tomography. Dev. Cell 57, 1132–1145.e5 (2022).

109. Sheng, X. et al. Structural insight into light harvesting for photosystem II in green algae. Nat Plants 5, 1320–1330 (2019).

110. Strauss, M., Hofhaus, G., Schröder, R. R. & Kühlbrandt, W. Dimer ribbons of ATP synthase shape the inner mitochondrial membrane. EMBO J. 27, 1154–1160 (2008).

111. Davies, K. M., Anselmi, C., Wittig, I., Faraldo-Gómez, J. D. & Kühlbrandt, W. Structure of the yeast F_1_F_o_-ATP synthase dimer and its role in shaping the mitochondrial cristae. Proc. Natl. Acad. Sci. U. S. A. 109, 13602–13607 (2012).

112. Buzzard, E. et al. The consequence of ATP synthase dimer angle on mitochondrial morphology studied by cryo-electron tomography. Biochem. J 481, 161–175 (01 2024).

113. Murphy, B. J. et al. Rotary substates of mitochondrial ATP synthase reveal the basis of flexible F1-Fo coupling. Science 364, (2019).

114. Dietrich, L., Agip, A.-N. A., Kunz, C., Schwarz, A. & Kühlbrandt, W. In situ structure and rotary states of mitochondrial ATP synthase in whole Polytomella cells. Science 385, 1086–1090 (2024).

115. Kühlbrandt, W. Structure and Mechanisms of F-Type ATP Synthases. Annu. Rev. Biochem. 88, 515–549 (2019).

116. Vázquez-Acevedo, M. et al. The mitochondrial ATP synthase of chlorophycean algae contains eight subunits of unknown origin involved in the formation of an atypical stator-stalk and in the dimerization of the complex. J. Bioenerg. Biomembr. 38, 271–282 (2006).

117. Gupte, S. R., et al. CryoViT: Efficient segmentation of cryogenic electron tomograms with vision foundation models. bioRxiv (2024) doi:10.1101/2024.06.26.600701.

118. Zhao, Y. et al. Training-free CryoET Tomogram Segmentation. (2024) doi:10.48550/ARXIV.2407.06833.

119. Huang, Q., Zhou, Y. & Bartesaghi, A. MiLoPYP: self-supervised molecular pattern mining and particle localization in situ. Nat. Methods (2024) doi:10.1038/s41592-024-02403-6.

120. Zeng, X. et al. High-throughput cryo-ET structural pattern mining by unsupervised deep iterative subtomogram clustering. Proc. Natl. Acad. Sci. U. S. A. 120, e2213149120 (2023).

121. Liu, G. et al. DeepETPicker: Fast and accurate 3D particle picking for cryo-electron tomography using weakly supervised deep learning. Nat. Commun. 15, (2024).

122. Genthe, E. et al. PickYOLO: Fast deep learning particle detector for annotation of cryo electron tomograms. J. Struct. Biol. 215, 107990 (2023).

123. Wagner, T. & Raunser, S. The evolution of SPHIRE-crYOLO particle picking and its application in automated cryo-EM processing workflows. *Commun*. Biol. 3, (2020).

124. Chen, M. & Ludtke, S. J. Deep learning-based mixed-dimensional Gaussian mixture model for characterizing variability in cryo-EM. Nat. Methods 18, 930–936 (2021).

125. Powell, B. M. & Davis, J. H. Learning structural heterogeneity from cryo-electron sub-tomograms with tomoDRGN. bioRxiv (2023) doi:10.1101/2023.05.31.542975.

126. Luo, Z., Wang, Q. & Ma, J. OPUS-TOMO: Deep learning framework for structural heterogeneity analysis in cryo-electron tomography. bioRxiv (2024) doi:10.1101/2024.06.30.601442.

127. Li, X. et al. MPicker: Visualizing and picking membrane proteins for cryo-electron tomography. Research Square (2024) doi:10.21203/rs.3.rs-4404303/v1.

128. Umen, J. G. & Goodenough, U. W. Control of cell division by a retinoblastoma protein homolog in Chlamydomonas. Genes Dev. 15, 1652–1661 (2001).

129. Khavnekar, S., Erdmann, P. S. & Wan, W. TOMOMAN: a software package for large-scale cryo-electron tomography data preprocessing, community data sharing and collaborative computing. J. Appl. Crystallogr. 57, 2010–2016 (2024).

130. Zheng, S. Q. et al. MotionCor2: anisotropic correction of beam-induced motion for improved cryo-electron microscopy. Nat. Methods 14, 331–332 (2017).

131. Khavnekar, S. & Wan, W. An approach for coherent periodogram averaging of tilt-series data for improved CTF estimation. bioRxiv (2024) doi:10.1101/2024.10.10.617684.

132. Rohou, A. & Grigorieff, N. CTFFIND4: Fast and accurate defocus estimation from electron micrographs. J. Struct. Biol. 192, 216–221 (2015).

133. Zheng, S. et al. AreTomo: An integrated software package for automated marker-free, motion-corrected cryo-electron tomographic alignment and reconstruction. J Struct Biol X 6, 100068 (2022).

134. Tegunov, D. & Cramer, P. Real-time cryo-electron microscopy data preprocessing with Warp. Nat. Methods 16, 1146–1152 (2019).

135. Zivanov, J. et al. A Bayesian approach to single-particle electron cryo-tomography in RELION-4.0. Elife 11, (2022).

136. Mastronarde, D. N. & Held, S. R. Automated tilt series alignment and tomographic reconstruction in IMOD. J. Struct. Biol. 197, 102–113 (2017).

137. Righetto, R. & LorenzLamm. CellArchLab/Slabify-et: V0.2.0. (Zenodo, 2024). doi:10.5281/ZENODO.13941082.

138. Milicevic, N., Jenner, L., Myasnikov, A., Yusupov, M. & Yusupova, G. mRNA reading frame maintenance during eukaryotic ribosome translocation. Nature 625, 393–400 (2024).

139. alisterburt, Tegunov, D., tegunovd & Wachsmuth-Melm, M. Warpem/Warp: V2.0.0dev31. (Zenodo, 2024). doi:10.5281/ZENODO.13982246.

140. Himes, B. & Grigorieff, N. Cryo-TEM simulations of amorphous radiation-sensitive samples using multislice wave propagation. IUCrJ 8, 943–953 (2021).

141. Tachiwana, H. et al. Structural basis of instability of the nucleosome containing a testis-specific histone variant, human H3T. Proc. Natl. Acad. Sci. U. S. A. 107, 10454–10459 (2010).

142. Meng, E. C. et al. UCSF ChimeraX: Tools for structure building and analysis. Protein Sci. 32, e4792 (2023).

143. Zivanov, J. et al. New tools for automated high-resolution cryo-EM structure determination in RELION-3. Elife 7, (2018).

144. Rosenthal, P. B. & Henderson, R. Optimal determination of particle orientation, absolute hand, and contrast loss in single-particle electron cryomicroscopy. J. Mol. Biol. 333, 721–745 (2003).

145. Rigort, A. et al. Automated segmentation of electron tomograms for a quantitative description of actin filament networks. J. Struct. Biol. 177, 135–144 (2012).

146. Chakraborty, S., Mahamid, J. & Baumeister, W. Cryoelectron Tomography Reveals Nanoscale Organization of the Cytoskeleton and Its Relation to Microtubule Curvature Inside Cells. Structure 28, 991–1003.e4 (2020).

147. Pettersen, E. F. et al. UCSF Chimera--a visualization system for exploratory research and analysis. J. Comput. Chem. 25, 1605–1612 (2004).

148. Qu, K. et al. Structure and architecture of immature and mature murine leukemia virus capsids. Proc. Natl. Acad. Sci. U. S. A. 115, E11751–E11760 (2018).

149. Sosa, H. & Chrétien, D. Relationship between moiré patterns, tubulin shape, and microtubule polarity. Cell Motil. Cytoskeleton 40, 38–43 (1998).

150. Ermel, U. H., Arghittu, S. M. & Frangakis, A. S. ArtiaX: An electron tomography toolbox for the interactive handling of sub-tomograms in UCSF ChimeraX. Protein Sci. 31, (2022).

151. Pettersen, E. F. et al. UCSF ChimeraX: Structure visualization for researchers, educators, and developers. Protein Sci. 30, 70–82 (2021).

152. Scheres, S. H. W. & Chen, S. Prevention of overfitting in cryo-EM structure determination. Nat. Methods 9, 853–854 (2012).

153. Pellegrino, S. et al. Structural insights into the role of diphthamide on elongation factor 2 in mRNA reading-frame maintenance. J. Mol. Biol. 430, 2677–2687 (2018).

154. Hall, S. R. The STAR file: a new format for electronic data transfer and archiving. J. Chem. Inf. Comput. Sci. 31, 326–333 (1991).

155. Diogo Righetto, R. EMPIAR-11830 dataset annotation: in situ cryo-electron tomography of Chlamydomonas reinhardtii. Zenodo 10.5281/ZENODO.13941456 (2024).

156. Gui, M. et al. Structures of radial spokes and associated complexes important for ciliary motility. Nat. Struct. Mol. Biol. 28, 29–37 (2021).

